# Modules in connectomes of phase-synchronization comprise anatomically contiguous, functionally related regions

**DOI:** 10.1101/2021.06.24.449415

**Authors:** N Williams, SH Wang, G Arnulfo, L Nobili, S Palva, JM Palva

## Abstract

Modules in brain functional connectomes are essential to balancing segregation and integration of neuronal activity. Connectomes are the complete set of pairwise connections between brain regions. Non-invasive Electroencephalography (EEG) and Magnetoencephalography (MEG) have been used to identify modules in connectomes of phase-synchronization. However, their resolution is suboptimal because of spurious phase-synchronization due to EEG volume conduction or MEG field spread. Here, we used invasive, intracerebral recordings from stereo-electroencephalography (SEEG, *N* = 67), to identify modules in connectomes of phase-synchronization. To generate SEEG-based group-level connectomes affected only minimally by volume conduction, we used submillimeter accurate localization of SEEG contacts and referenced electrode contacts in cortical grey matter to their closest contacts in white matter. Combining community detection methods with consensus clustering, we found that the connectomes of phase-synchronization were characterized by distinct and stable modules at multiple spatial scales, across frequencies from 3 to 320 Hz. These modules were highly similar within canonical frequency bands. Unlike the distributed brain systems identified with functional Magnetic Resonance Imaging (fMRI), modules up to the high-gamma frequency band comprised only anatomically contiguous regions. Notably, the identified modules comprised cortical regions involved in shared repertoires of sensorimotor and cognitive functions including memory, language and attention. These results suggest that the identified modules represent functionally specialised brain systems, which only partially overlap with the brain systems reported with fMRI. Hence, these modules might regulate the balance between functional segregation and functional integration through phase-synchronization.

**Highlights:**

- Large-cohort SEEG used for phase-synchronization connectomics
- Connectomes of phase-synchronization possess distinct and stable modules
- Modules in connectomes are highly similar within canonical frequency bands
- Modules in connectomes comprise anatomically contiguous regions
- Modules in connectomes comprise functionally related regions

## 1. Introduction

Structural and functional connectomes obtained from Magnetic Resonance Imaging (MRI) possess a modular organization (Meunier et al. (2009), Power et al. (2011), Doucet et al. (2011)). Connectomes are the complete set of connections between brain regions. Modules are sets of strongly interconnected brain regions. Modules identified in resting-state fMRI comprise regions that have also been observed to be concurrently active during task processing and have been found to delineate functional systems for executive, attentional, sensory, and motor processing (Beckmann et al. (2005), Smith et al. (2009), Yeo et al. (2011), Cole et al. (2014)). The anatomical structure of resting-state modules in fMRI connectomes has been found to be reproducible and similarly observable with different approaches such as community detection (Valencia et al. (2009), Power et al. (2011)) and clustering (Benjaminsson et al. (2010), Yeo et al. (2011), Lee et al. (2012)). Moreover, the balance between segregated information processing in modules (Wig (2017)) and integrated information processing via inter-modular connections, is essential to brain functioning (Tononi et al. (1994), Tononi et al. (1998), Deco et al. (2015)).

The relationship of fMRI functional connectivity to underlying electrophysiological connectivity is complex and not attributable to any single form of neuronal activity or coupling (Kucyi et al. (2018), Shafiei et al. (2022)). Electrophysiological measurements of macro-scale neuronal activity with Magneto-(MEG) and Electroencephalography (EEG) reveal band-limited neuronal oscillations in multiple frequencies, whose inter-regional coupling is observable as synchronization between oscillation phases and correlations between oscillation amplitude envelopes (Palva et al. (2005), Fell & Axmacher (2011), Brookes et al. (2011), Palva & Palva (2012), Engel et al. (2013)). Amplitude correlations reflect, *e.g.*, co-modulation in neuronal excitability (Vanhatalo et al. (2004), Schroeder & Lakatos (2009), Engel et al. (2013)) while phase-synchronization implies spike-time relationships of neuronal activity and may regulate inter-regional neuronal communication (Fries (2015), Bastos et al. (2015)). Large-scale networks of phase-synchronization are proposed to support the coordination, regulation, and integration of neuronal processing in cognitive functions, both in frequencies up to 130 Hz (Varela (2001), Palva et al. (2005), Uhlhaas et al. (2010), Kitzbichler et al. (2011), Palva & Palva (2012)), and in frequencies higher than 130 Hz, *i.e.,* high-frequency oscillations (HFO) (Vaz et al. (2019), Arnulfo et al. (2020)).

In the light of such putative mechanistic roles for phase-synchronization in cognitive functions, a modular architecture and inter-modular coupling in connectomes of phase-synchronization during resting-state would establish a baseline to support corresponding demands for functional segregation and integration during cognitive operations (Smith et al. (2009), Spadone et al. (2015)). An MEG study investigated modules in connectomes of phase-synchronization and amplitude correlation using source-reconstructed resting-state data (Zhigalov et al. (2017)). Both connectomes of amplitude correlation and phase-synchronization comprised distinct modules in frontal regions, sensorimotor regions and occipital regions, particularly in the alpha (8–14 Hz) and beta (14–30 Hz) frequency bands. Another MEG study used source-reconstructed resting-state data to identify module-like structures in connectomes of inter-regional coherence (Vidaurre et al. (2018)), a connectivity measure influenced by phase-synchronization. The connectomes included module-like structures comprising frontal regions, sensorimotor regions and occipital regions across delta/theta (1–8 Hz), alpha (8–14 Hz) and beta (14–30 Hz) frequencies. However, the accuracy of modules identified in MEG/EEG connectomes is compromised by the intrinsic resolution limitations of these methods, including artificial and spurious false positive observations with bivariate connectivity measures arising from source leakage (Palva & Palva (2012), Palva et. al (2018)) as well as false negatives due to linear-mixing insensitive measures that ignore also true near-zero-lag phase-synchronization (Vinck et al. (2011), Brookes et al. (2012), Palva & Palva (2012)). On the other hand, low-resolution (< 35 parcels /hemisphere) cortical parcellations, which are needed when spurious connections are eliminated by multivariate leakage correction (Colclough et al. (2015)), may be too coarse to identify fine-grained cortical network structures such as modules.

In this study, we pooled resting-state stereo-EEG (SEEG) recordings data from a large cohort (*N* = 67) to accurately estimate connectomes of phase-synchronization. In contrast to the centimetre-scale, insight yielded by MEG, SEEG provides a millimetre range, meso-scale measurement of human cortical local field potentials (LFPs) (Parvizi & Kastner (2018), Zhigalov et al. (2015), Zhigalov et al. (2017)). We used submillimetre-accurate anatomical localization of SEEG electrode contacts to brain regions (Narizzano et al. (2017), Arnulfo et al. (2015b)) and referenced each gray-matter contact to its closest white-matter contact (Arnulfo et al. (2015a)), which yielded polarity-correct measurements of local cortical activity without the phase distortion potentially arising with conventional bipolar referencing. This enabled the estimation of a large proportion of connections in the connectome while adequately controlling for volume conduction so that also near zero-lag phase-synchronization was measurable (Arnulfo et al. (2015a)). Finally, we combined community detection with consensus clustering (Williams et al. (2019)) to identify modules in connectomes of phase-synchronization in a manner that is robust against unsampled connections.

We found that connectomes of phase-synchronization exhibited modular organization at multiple spatial scales, throughout the studied range of frequencies from 3 to 320 Hz. These modules were highly similar within canonical frequency bands and comprised anatomically contiguous regions up to the high-gamma frequency band (80-113 Hz). Finally, we used Neurosynth meta-analysis decoding (Yarkoni et al. (2011)) to reveal that the observed modules comprised cortical regions exhibiting shared cognitive functions, suggesting that these modules correspond to brain systems with specific functional roles. Hence, the modules identified might serve the regulation of balance between segregation and integration of neuronal activity through phase-synchronization.

## 2. Materials & Methods

### 2.1 Analysis pipeline to identify modules in connectomes of phase-synchronization

We combined pre-surgical SEEG recordings from epileptic patients with state-of-the-art methods, to identify modules in connectomes of phase-synchronization. We recorded resting-state LFP data from each patient using a common reference in white matter, distant from the putative epileptogenic zone. We re-referenced the LFP activity of each grey-matter SEEG contact to its closest white-matter contact, which we have demonstrated to preserve undistorted phase reconstruction while minimising volume conduction (Arnulfo et al. (2015a)). We filtered the recorded LFP data using 18 narrow-band Finite Impulse Response (FIR) filters (Figure 1A) from 2.5 Hz up to 350 Hz with line-noise suppressed using band-stop filters at 50 Hz and harmonics. Next, we estimated the strength of phase-synchronization between every pair of SEEG contacts, for each frequency, using Phase Locking Value (Figure 1B). We assigned each SEEG contact to a brain region, by first identifying the position of each contact from a post-implant CT volume, and using co-registered pre-implant MRI scans to assign each contact to one of 148 regions in the Destrieux brain atlas (Destrieux et al. (2010)) with FreeSurfer (http://freesurfer.net/). We identified the position of each SEEG contact by using planned entry and termination points of SEEG shafts to initialize the shaft axis, and used constraints of inter-contact distance and axis deviation to locate each SEEG contact along the shaft axis (Arnulfo et al. (2015b)). We then estimated group-level connectomes by averaging for each region-pair, the corresponding contact-contact PLVs across subjects (Figure 1C). We analyzed the left and right hemispheres separately (Figure 1D) and identified modules with Louvain community detection (Blondel et al. (2008)) combined with consensus clustering (Williams et al. (2019)) (Figure 1E). Finally, we visualised the identified modules on anatomical brain surfaces (Figure 1F).

**Figure 1.**
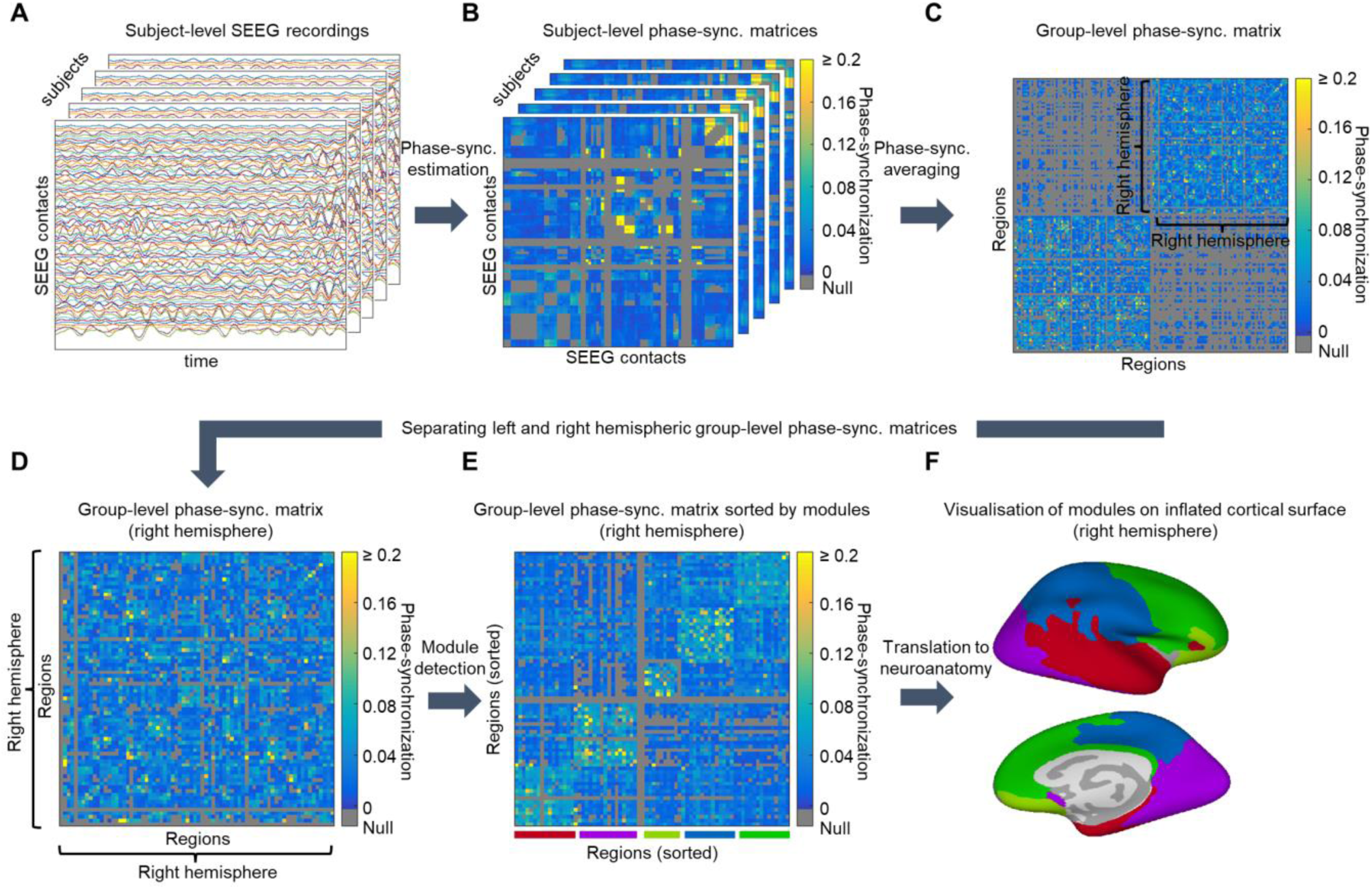
Modules in connectomes of phase-synchronization estimated by pooling data across subjects. **A.** Band-pass filtered data (center frequency=14 Hz) for example group of subjects. **B.** Subject-evel matrices of phase-synchronization between SEEG contacts, for example group of subjects. **C.** Group-evel matrix of phase-synchronization between brain regions. Matrix ordered to show left-(bottom left), ight-(top right) and inter-hemispheric connections (top left and bottom right) respectively. Non-estimable connections are gray. **D.** Group-level matrix of phase-synchronization between right-hemispheric regions. **E.** Sorted group-level matrix of phase-synchronization between right-hemispheric regions, sorting based n results of community detection to identify modules. **F.** Colour-coded modules for lateral (top) and medial bottom) views of right-hemispheric inflated cortical surface.

### 2.2 Data acquisition

We recorded SEEG data from 67 participants affected by drug-resistant focal epilepsy and undergoing pre-surgical clinical assessment. For each participant, we inserted 17 ± 3 (mean ± SD) SEEG shafts into the brain, with anatomical positions varying by surgical requirements. Each shaft had between 8 and 15 platinum-iridium contacts, each contact being 2 mm long and 0.8 mm thick, with inter-contact distance of 1.5 mm (DIXI medical, Besancon, France). We acquired 10 minutes eyes-closed resting-state activity from each participant, via a 192-channel SEEG amplifier system (Nihon Kohden Neurofax-110) at a sampling frequency of 1 kHz. We obtained written informed consent from participants prior to recordings. We obtained ethics approval for the study from Niguarda “Ca’ Granda” Hospital, Milan, and we performed the study according to WMA Declaration of Helsinki – Ethical Principles for Medical Research Involving Human Subjects.

### 2.3 Pre-processing

We performed re-referencing, filtering and artefact removal of the SEEG data, before estimating the connectome of phase-synchronization. We originally recorded data from all contacts with a monopolar referencing scheme. We subsequently re-referenced activity from each gray-matter contact to the nearest white matter contact as identified by GMPI (gray matter proximity index). We have previously demonstrated the utility of this referencing scheme in studying phase-synchronization, since phase relationships between contacts are well preserved (Arnulfo et al. (2015a)). We only analysed activity from gray-matter contacts after re-referencing. We filtered activity from each gray-matter contact using FIR filters (equiripples 1% of maximal band-pass ripples) into 18 frequency bands, with center frequencies (*F_c_*) ranging from 3 to 320 Hz (excluding 50 Hz line-noise and harmonics). We used log-spaced center frequencies of 3 Hz, 4 Hz, 5 Hz, 7 Hz, 10 Hz, 14 Hz, 20 Hz, 28 Hz, 40 Hz, 57 Hz, 80 Hz, 113 Hz, 135 Hz, 160 Hz, 190 Hz, 226 Hz, 269 Hz and 320 Hz. We used a relative bandwidth approach for filter banks such that pass band (*W_p_*) and stop band (*W_s_*) were defined 0.5 × *F_c_* and 2 × *F_c_*, respectively for low and high-pass filters, producing log-increasing spectral window widths. The choice of log-spaced center frequencies followed the experimentally observed center frequencies of brain oscillations (Penttonen & Buzsáki (2003)). The log-increasing window widths afforded fine spectral resolution at lower frequencies, avoiding confounding instantaneous phases of multiple frequency components at lower frequencies (Lopes da Silva (2013)). Simultaneously, this choice also provided fine temporal resolution at higher frequencies, enabling accurately estimating the instantaneous phase of the known-to-be-short-lived oscillations at higher frequencies (Lundquist et al. (2018)). We applied the Hilbert transform to the FIR-filtered signal to return the analytic signal, from which angle we extracted the instantaneous phase. Before estimating phase-synchronization, we excluded select 500 ms windows containing Inter-Ictal Epileptic (IIE) events, to counteract any spurious phase-synchronization due to filtering artefacts around the epileptic spikes. We defined IIE as at least 10% of SEEG contacts narrow-band time series demonstrating abnormal, concurrent sharp peaks in more than half the 18 frequencies. To identify such periods, we searched for “spiky” periods in amplitude envelopes of each SEEG contact. We tagged a 500 ms window as “spiky” if any of its samples were 5 standard deviations higher than mean amplitude of the contact.

### 2.4 Connectome estimation

We pooled estimates of phase-synchronization between SEEG contacts to obtain the group-level inter-regional connectome of phase-synchronization. We measured phase-synchronization between SEEG contacts with Phase Locking Value (Lachaux et al. 1999):

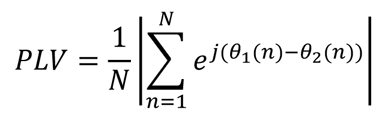

where *θ_1_*(*n*) and *θ_2_(n)* are instantaneous phases from a pair of SEEG contacts at sample *n*, with *N* being the total number of samples. We estimated the group-level phase-synchronization between a pair of brain regions as the average PLV over all subjects, of all SEEG contact-pairs traversing that pair of brain regions. This procedure furnished accurate estimates of group-level phase-synchronization, since it computes a weighted average of phase-synchronization across subjects, wherein subjects contributing higher number of PLV values are assigned a higher weight in the group-level estimate. The alternative procedure of first averaging all PLV values for a pair of brain regions for each subject separately, before averaging these subject-level PLV estimates, would assign equal weight to each subject in the group-level estimate despite some subjects contributing higher number of PLV values to the estimate. We estimated the connectome of phase-synchronization as the group-level phase-synchronization between every pair of 148 regions in the Destrieux brain atlas for which we had at least one SEEG contact-pair. We then thresholded the connectome by retaining the estimated strengths of only the top 20 percentile of connections, setting all others to 0. We performed this thresholding as a means of emphasising the topological organisation of the connectome (Rubinov & Sporns (2010)). We also determined the robustness of our results to the specific choice of percentile threshold, by also identifying modules on connectomes thresholded by retaining the strengths of the top 10 and top 30 percentile of connections.

Since we did not have complete recording coverage of the brain with SEEG, we had insufficient data to estimate phase-synchronization of all connections in the group-level connectome. Rather, we had sufficient coverage with SEEG, to estimate phase-synchronization of 47.2% of connections in the group-level connectome. Many of these connections were intra-hemispheric - we estimated phase-synchronization of 68% of connections between just left-hemispheric regions, and of 80% of connections between just right-hemispheric regions. Hence, we separately identified modules in the connectome of just left-hemispheric regions and in the connectome of just right-hemispheric regions.

We excluded selected contact-pairs from the connectome estimation due to potential artefacts, as per the below criteria. We excluded contact-pairs involving SEEG contacts marked by clinical experts as falling within the epileptogenic or seizure propagation regions. We performed this step after we removed 500 ms windows containing IIE, as described above (Section 2.3). Further, we excluded contact-pairs whose respective SEEG contacts were less than 20 mm apart and those with the same white-matter reference, both to reduce the effect of volume conduction. We have described these steps in further detail, in recent work using the same SEEG dataset (Arnulfo et al. (2020)).

### 2.5 Analysing the connectome of phase-synchronization

#### 2.5.1 Identifying modules in connectomes of phase-synchronization

We used Louvain community detection (Reichardt & Bornholdt (2006), Blondel et al. (2008), Ronhovde & Nussinov (2009), Sun et al. (2009)) combined with consensus clustering (Lancichinetti & Fortunato (2012)) to identify modules in the connectome of phase-synchronization. Modules are sets of strongly interconnected nodes in a network. The Louvain community detection method iteratively identifies a partition of network nodes into modules, such that ‘modularity’ of the partition is maximised. The ‘modularity’ objective function that is maximised, quantifies the extent to which the network comprises non-overlapping modules compared to a null model of an equivalent network that would be expected by chance (Blondel et al. (2008)). We chose the Louvain method due to its superior performance in accurately identifying network modules compared to alternative community detection methods (Lancichinetti & Fortunato (2009)), and its superior performance, when combined with consensus clustering, in recovering modules in incomplete brain networks (Williams et al. (2019)). We used the implementation of the Louvain method in Brain Connectivity Toolbox (Rubinov & Sporns (2010)). We applied the Louvain method to left and right hemispheric regions separately, since the low number of inter-hemispheric connections might confound the identification of modules. To identify modules while accounting for missing values in the group-level connectome matrix, we first generated 5000 variants of the connectome wherein we replaced each missing value with a randomly sampled (with replacement) existing value from the group-level connectome. Replacing missing values with existing values from the group-level connectome generates complete connectomes with the same distribution of phase-synchronization strengths as the original incomplete connectome. We applied Louvain community detection to identify modules on each of these 5000 complete matrices. We identified modules at a range of spatial scales by setting the γ input parameter of the Louvain method from 0.8 to 5, in intervals of 0.1. For each γ value, we combined the module assignments of the 5000 connectome variants to obtain a consensus module assignment. We performed this step by first generating matrix representations of each module assignment, with number of matrix rows and columns being the number of regions. We set each element in the matrix to 1 or 0 depending respectively on whether that pair of regions were in the same module or not. We then obtained a consensus matrix by averaging the 5000 matrix representations and obtained a consensus module assignment by applying the Louvain method to this consensus matrix. We have demonstrated this consensus clustering approach is superior to other approaches to identify modules in incomplete human brain networks (Williams et al. (2019)). We applied this procedure to identify modules at each frequency, for left and right hemispheres separately.

#### 2.5.2. Determining statistical significance of modular organization

We determined statistical significance of modular organization by comparing modularity of connectomes against modularity of corresponding randomized connectomes. Modularity is high when the connectome is divided into internally dense modules. We compared modularity of the original connectomes to their corresponding randomized connectomes, with the following steps 1.) We estimated modularity of the original connectome using Louvain community detection in combination with consensus clustering, for γ values (spatial scales) from 0.8 to 5 (see Section 2.5.1). Modularity is the objective maximised by the Louvain method. We used 100 variants of the original connectome for the consensus clustering step. 2.) We standardised the modularity values of the original connectomes by *z*-scoring the estimated modularity at each γ value against a null distribution of 100 modularity values generated by identifying modules on randomly permuted (without replacement) versions of the original connectome. We identified modules for each of these randomized connectomes with the same procedure as we used to identify modules on the original connectome. We estimated *z*-scored modularity for connectomes at each frequency, for left and right hemispheres separately. 3.) We determined the statistical significance of the estimated modularity values of the original connectome by converting the *z*-scores to *p*-values assuming a Gaussian distribution and used False Discovery Rate (FDR) thresholding (Benjamini & Hochberg (1995)) to correct for multiple comparisons across all combinations of γ and frequency. We considered FDR-corrected *p* < 0.05 to indicate statistically significant modular organization of the original connectome at a given γ and frequency. We performed FDR thresholding separately for connectomes of each hemisphere.

#### 2.5.3 Determining statistical significance of percentage of stable regions

We determined the statistical significance of percentage of stable regions using a permutation-based test to assess the stability of module assignment of each brain region, and a second permutation-based test to assess if the percentage of stable regions is higher than expected by chance. We performed the following steps 1.) We constructed 100 bootstrapped connectomes with the same procedure as for the original connectomes (Section 2.4), each from a cohort of 67 randomly resampled (with replacement) subjects from the original cohort. 2.) We considered the module assignment of a brain region to be stable if it was assigned to the same module in the original connectome, as it was assigned to across the 100 bootstrapped connectomes. Hence, we quantified the stability of module assignment of a region as the mean correspondence in its module affiliation in the original connectome, to module affiliations of the same region across the 100 bootstrapped connectomes. For a given brain region, we specified module affiliation as a vector of ‘1’ and ‘0’s, depending respectively on whether each other brain region was or was not assigned to the same module, and we estimated the correspondence between module affiliations by the proportion of common ‘1’s and ‘0’s. Highly stable assignment of modules for a given brain region, were reflected in mean correspondences in module affiliation close to 1, for that brain region. 3.) We counted the stability of module assignment of a brain region as statistically significant if it exceeded the 95-percentile value of the null distribution of stability values for that brain region. We estimated the null distribution of stability values as the mean stability values when comparing module affiliation with the original connectome against 100 randomly permuted (without replacement) module affiliation vectors of each of the bootstrapped connectomes. Hence, we had 100 samples in the null distribution of stability values for each brain region, one for each bootstrapped connectome. 4.) We next estimated the percentage of brain regions for each combination of spatial scales or γ values (from 0.8 to 5) and frequencies, for left and right hemispheres separately. 5.) Finally, we determined the statistical significance of the percentage of stable regions, by *z*-scoring it against the percentage of regions expected to be stable by chance across the 100 bootstrapped connectomes. We estimated these chance percentages for each bootstrapped connectome, as the percentage of brain regions whose null stability values exceeded the 95-percentile value of the null distribution of stability values for that bootstrapped connectome. We then converted the *z*-scores to *p*-values assuming a Gaussian distribution and used False Discovery Rate (FDR) thresholding to correct for multiple comparisons due to testing across every combination of γ and frequency. We considered FDR-corrected *p* < 0.05 to indicate statistically significant percentage of stable regions.

#### 2.5.4. Grouping frequencies by similarity of modules

We used multi-slice community detection (Mucha et al. (2010)) to identify groups of frequencies with similar modules, simultaneously for both left and right hemispheres. First, we generated matrices of module similarity between each pair of frequencies, separately for left and right hemispheres. We estimated similarity between module assignments by first generating matrix representations of module assignments at each frequency. The number of rows and columns of these matrices were equal to the number of brain regions, each element being set to 1 or 0 depending respectively on whether the corresponding pair of brain regions were in the same module or not. We measured similarity between module assignments using partition similarity (Ben-Hur et al. (2002)):

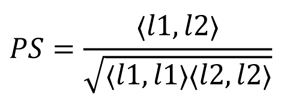

where 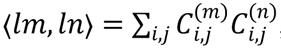, *i.e.,* the dot product between matrix representations of the module assignments for frequencies *m* and *n*. Note that this measure of partition similarity effectively extends the measure in Section 2.5.3, which compares the module assignments of single brain regions (in relation to a set of brain regions), to the case of comparing the module assignments of a set of brain regions. We obtained matrices of partition similarity for each γ value (spatial scale) from 0.8 to 5. We then estimated a weighted average of these matrices across the γ dimension, to yield a matrix indicating similarity of modules between frequencies that was consistent across spatial scales. We assigned weights to the matrix at each γ value, as the number of frequencies with statistically significant modular organisation at that γ value. Note however, that we also compared the frequency groupings we obtained when applying these weights, to frequency groupings we obtained when applying equal (unit) weights to the module similarity matrices at all γ values.

We entered the left and right hemispheric matrices of module similarity into a multi-slice community detection procedure (Mucha et al. (2010)), to identify groups of frequencies with similar modules for both hemispheres. Multi-slice community detection is a principled generalisation of modularity maximisation community detection methods, *e.g.,* Louvain, to multiple slices. It does so by formulating the null model for community structure across multiple slices, in terms of the stability of communities under Laplacian dynamics (Mucha et al. (2010)). Multi-slice community detection has previously been applied to study dynamic reconfiguration of human brain networks in learning (Bassett et al. (2011)), and to relate modules in human brain networks identified for different cognitive tasks (Cole et al. (2014)).

The method has two input parameters, γ_multislice_ and ω. γ_multislice_ represents the spatial scale (just as with γ for the Louvain method), while ω represents the dependence between communities across the different slices. In our context, γ_multislice_ influences the number of identified groups of frequencies while ω controls the dependence between the identified groups of left and right hemispheres. To select values for these parameters, we first estimated modularity values for each combination of γ_multislice_ = 1 – 1.5 (intervals of 0.05) and ω = 0.1 – 1 (intervals of 0.1). Then, we generated a null distribution of modularity values by applying the method to identically randomly resampled (without replacement) left and right hemispheric matrices of module similarity. We *z*-scored the original modularity values against the null distribution and converted them to *p*-values assuming a Gaussian distribution. Finally, we inspected frequency groups for selected combinations of γ_multislice_ and ω with FDR-thresholded *p* < 0.05.

#### 2.5.5 Identifying modules across multiple frequencies or spatial scales

We used a consensus clustering approach (Section 2.5.1) to identify a single set of modules across frequencies and spatial scales. To do this, we first generated matrix representations of modules at individual frequencies, at each γ value (spatial scale) from 0.8 to 5, for left and right hemispheres separately. Matrix representations have number of rows and columns equal to the number of brain regions, each element in the matrix is 1 or 0 depending respectively on whether the corresponding pair of regions are in the same module or not. We then averaged the matrix representations, first across all frequencies and then across all spatial scales, for left and right hemispheres separately. Finally, we applied multi-slice community detection to the averaged matrices of left and right hemispheres, to identify eight modules representing sets of regions assigned to the same module across frequencies and spatial scales, for both left and right hemispheres. The rationale for identifying these consensus modules was to relate these modules to their putative fMRI counterparts, Resting State Networks (RSNs). Hence, we fixed γ_multislice_ to 1.6 to return eight modules, while we set ω = 1 to constrain the modules to be bilaterally symmetric – RSNs are typically reported as between seven and ten bilaterally symmetric modules (*e.g.,* in Yeo et al. (2011)).

### 2.6 Inferring whether regions in a module are functionally related

We combined Neurosynth meta-analyses decoding (Yarkoni et al. (2011)) with comparison to surrogate modules, to assign putative functional roles to each module. We used Neurosynth decoding to find terms related to perception, cognition and behaviour selectively associated to the centroid co-ordinates of each brain region, based on a large database of fMRI studies. Then, we aggregated the terms associated with regions in each module and compared the occurrence frequencies of these terms to those of equally sized surrogate modules which were constrained to comprise anatomically proximal and bilaterally symmetric brain regions. Hence, we determined terms that were common to regions in a module, even after accounting for the anatomical proximity of its regions. We *z*-scored the occurrence frequency of each term in a module against corresponding frequencies of the surrogate modules. We converted these *z*-scores to *p*-values assuming a Gaussian distribution and FDR-thresholded at *p* < 0.05, to reveal those terms selectively associated to each module.

We inferred the putative functional role of each module by the set of terms it was selectively associated to. We also performed a post-hoc analysis to verify the functional specificity of each module. To do this, we generated an 8 × 8 ‘confusion matrix’ of percentages of selectively associated terms of each module distributed across the eight cognitive functions assigned to the modules. High values along the diagonal would reflect high functional specificity, *i.e.,* that the terms of each module were largely confined to a single cognitive function. We compared these percentages against the percentages of all terms related to a module, not just those selectively associated to each module. We expected these sets of all terms of each module to be distributed across diverse cognitive functions.

### 2.7 Assessing robustness of modules identified

We assessed robustness of modules identified, to a range of potential confounds. First, we assessed the robustness of modules identified to the specific set of SEEG contact-pairs used to generate the group-level connectomes of phase-synchronization. To do this, we identified and compared modules identified from split connectomes at γ = 2, each of the split connectomes being generated by combining different sets of SEEG contact-pairs. To generate a split connectome, we estimated strength of each connection from a randomly selected sample of half the SEEG contact-pairs used to estimate strength of each estimated connection in the original connectome. We estimated the same connection in the other split connectome with the other half of SEEG contact-pairs used to estimate strength of that connection in the original connectome. Next, we assessed the robustness of the modules to the community detection method used to identify the modules. To do this, we compared the original modules obtained with Louvain community detection at γ = 2, against modules obtained with Infomap community detection (Rosvall & Bergstrom (2008)). Network density influences the number of modules with Infomap - we set the network density to 10% since this value yielded interpretable modules in previous work (Williams et al. (2019)). Further, we assessed the robustness of our results when modules were identified on binarized rather than weighted connectomes, when modules were identified by retaining the top 10 and 30 percentile group-level connections rather than top 20 percentile, and when modules were identified on connectomes generated with a criterion of at least 5 and 10 SEEG contact-pairs required to estimate an inter-regional group-level connection, rather than at least 1. Finally, we investigated if identifying modules is confounded by amplitude of oscillations from individual nodes in a network. To do this, we compared modules of the 67 subject-level networks of phase-synchronization before and after removing amplitude-related differences in functional connection strengths, at each of the 18 frequencies, at six spatial scales (γ = 1, 1.8, 2.6, 3.4, 4.2 and 5). We removed amplitude-related differences by relating the strengths of each functional connection to average amplitude of corresponding node-pairs via linear regression, and recovering the residuals. We compared modules identified before and after removing amplitude-related differences, with the partition similarity measure (Section 2.5.4).

We have made available MATLAB code to perform each stage of the analyses, via our GitHub repository (https://github.com/nitinwilliams/eeg_meg_analysis/tree/master/FC_modules).

## 3. Results

### 3.1 Whole-brain coverage achieved by broad spatial sampling of SEEG contacts

We quantified the sampling of brain regions and inter-regional connections (Arnulfo et al. (2020)) by the percentage of brain regions and region-pairs in Destrieux brain atlas (Destrieux et al. (2010)) containing at least one gray-matter SEEG contact or an inter-regional SEEG-contact-pair across subjects, respectively. The cohort sampled with at least one SEEG contact 97% of brain regions (143 of 148) in the Destrieux atlas (Figure 2A). The SEEG contacts were sampled more densely on the right (*N* = 45 ± 38, mean ± standard deviation, range 0–123, contacts per subject) than the left (32 ± 41, 0–128, contacts per subject) hemisphere. This yielded a coverage of 68% of left-hemispheric, 80% of right-hemispheric connections and 20% of inter-hemispheric connections (Figure 1B). We also estimated the numbers of SEEG contacts across subjects in each of the Yeo functional systems (Yeo et al. (2011), Figure 1C) and found them densely sampled, with > 100 contacts in each functional system (Figure 1D).

**Figure 2.**
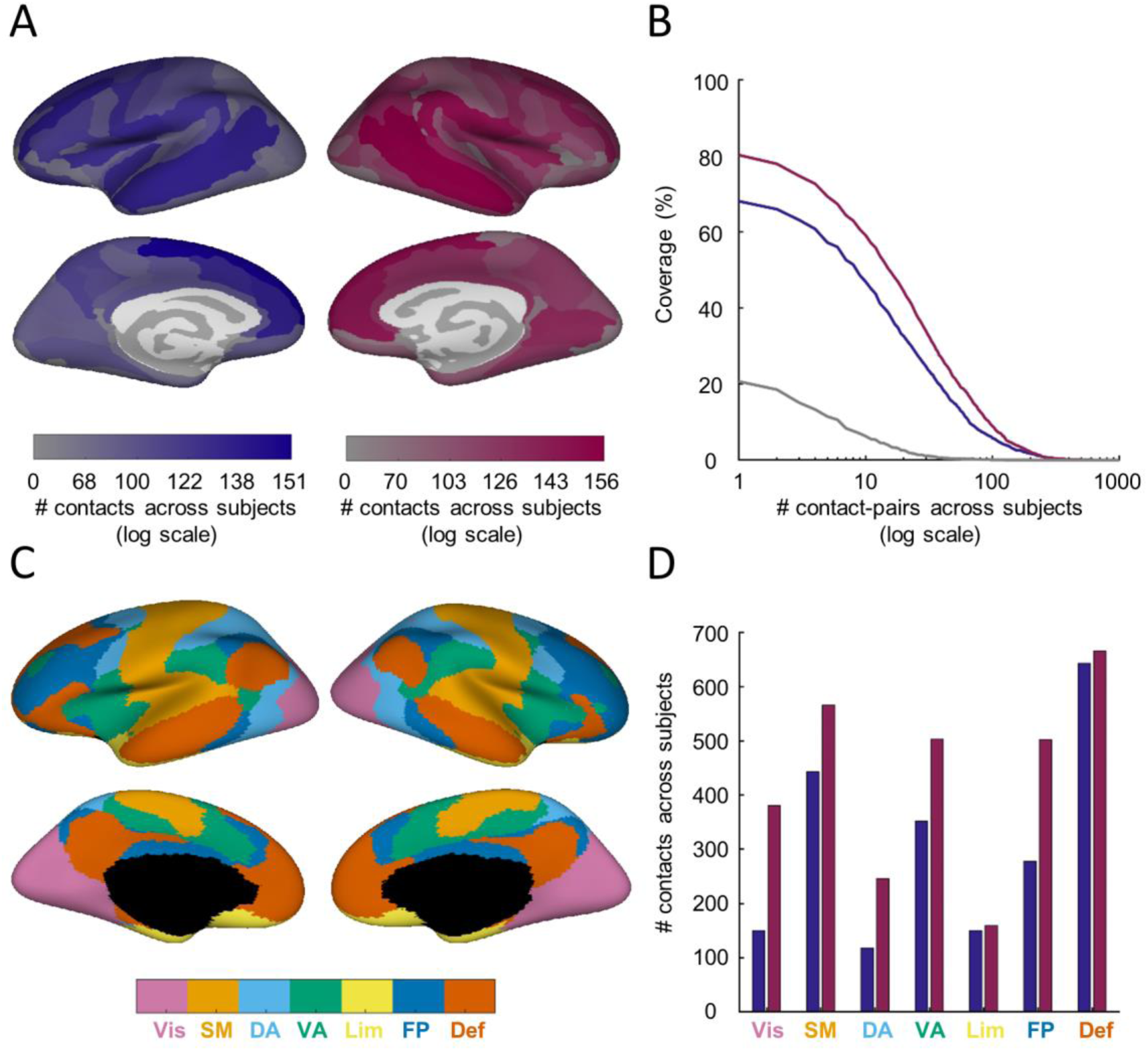
Whole-brain coverage achieved by placement of SEEG contacts. **A.** Number of SEEG contacts across subjects, in each brain region, for left (dark blue) and right (dark red) hemispheres, from lateral (top) and medial (bottom) views. **B.** Coverage of left-hemispheric (dark blue), right-hemispheric (dark red) and inter-hemispheric (gray) connections for a range of minimum number of SEEG contact-pairs across subjects. **C.** 7 Yeo systems from lateral (top) and medial (bottom) views. VIS = Visual, SM = Sensorimotor, DA = Dorsal Attention, VA = Ventral Attention, Lim = Limbic, FP = Fronto-parietal and Def = Default Mode. **D.** Number of SEEG contacts across subjects, in each of 7 Yeo systems, for left (dark blue) and right (dark red) hemispheres.

Within each hemisphere, we further investigated the coverage of inter-regional connections with respect to distance, provided by the SEEG recordings. Coverage of inter-regional connections both between proximal and between distant brain regions would allow the community detection method to identify modules comprising both proximal and distant brain regions, while coverage of connections between only proximal brain regions would limit the community detection method to identifying modules comprising only proximal brain regions. We expected coverage of connections both between proximal and between distant brain regions due to the large cohort of subjects we used (*N* = 67), with recordings from 17 ± 3 (mean ± SD) SEEG shafts from each subject. To assess the coverage with respect to distance, we 1.) determined the percentage of connections sampled by at least one SEEG contact-pair, for four distance categories: very short (< 30 mm), short (30–60 mm), medium (60–90 mm) and long (> 90 mm), for both left and right hemispheres, and 2.) determined the percentage of connections sampled by at least one SEEG contact-pair between regions in every pairwise combination of the following functional subdivisions: frontal, parietal, temporal, occipital, limbic and insula. We found that our SEEG recordings sampled inter-regional connections at all distance categories, for both hemispheres (Supplementary material, Figure S1A). Short-distance connections were sampled most densely, with 87% for left hemisphere and 92% for right hemisphere, but we also sampled 25% of long-distance connections for left hemisphere and 47% of long-distance connections for right hemisphere. Crucially, we found the SEEG recordings allowed dense sampling of inter-regional connections between standard functional subdivisions, *i.e.* frontal, parietal, temporal, occipital, limbic and insular cortices (Supplementary material, Figure S1B): between 48% and 100% of connections between regions in pairs of functional subdivisions were sampled for left hemisphere, and between 55% and 100% of connections were sampled for right hemisphere. Hence, the SEEG recordings allowed sampling intra-hemispheric connections both between proximal and between distant brain regions, including between regions in different functional subdivisions.

We identified modules on thresholded connectomes, wherein we retained the strengths of the top 20 percentile strongest connections, setting all others to 0. To check the sampling statistics, we investigated the relationship between the percentage of supra-threshold connections and connection distance. We found that the percentage of supra-threshold connections was higher for short-distance than long-distance connections. However, we did find several connections between spatially distant brain regions, including between regions in different functional subdivisions, *i.e.* frontal, parietal, temporal, occipital, limbic and insular cortices (Supplementary material, see Supplementary Text and Figure S2 for details).

### 3.2 Connectomes of phase-synchronization are characterized by distinct and stable modules at multiple spatial scales

We combined Louvain community detection with consensus clustering to identify modules in connectomes of phase-synchronization. The presence of distinct and stable modules would suggest that these modules operate as functional systems within the connectome. Hence, we determined the distinctness and stability of the identified modules. We performed this investigation at multiple spatial scales in order to avoid missing modules due to the resolution limit imposed by identifying modules at a single spatial scale (Sporns & Betzel (2016)). We used Louvain community detection with a range of the γ parameter from 0.8 to 5 to identify modules at multiple spatial scales. The numbers of modules varied from 1 to 18 across the range of spatial scales and filter center frequencies (Figure 3A). We used permutation-based methods to assess the distinctness and stability of the identified modules. To assess stability of the identified modules, we determined if the percentage of brain regions consistently assigned to the same module across bootstrapped versions (*N* = 100) of the original connectome, was more than would be expected by chance. To assess distinctness of the identified modules, we assessed if modularity of the original connectome was higher than modularity of randomized versions of the original connectome (*N* = 100). Modularity is high when the connectome divides into internally dense modules. We observed that across a wide range of spatial scales and frequencies, 12.2–100% cortical regions had stable module assignments, yielding statistically significant percentages of stable regions at multiple spatial scales (*p* < 0.05, FDR-corrected, permutation test) (Figure 3B). Further, the connectomes had statistically significantly distinct modular organization (*p* < 0.05, FDR-corrected, permutation test) at multiple spatial scales throughout the studied frequency range (Figure 3C). Connectomes in beta frequency band (14-20 Hz) exhibited the widest range of spatial scales for which modules were statistically significantly distinct. The distinctness and stability of the modules, across a range of spatial scales, suggests that modules of different sizes operate as functional systems within the connectome.

**Figure 3.**
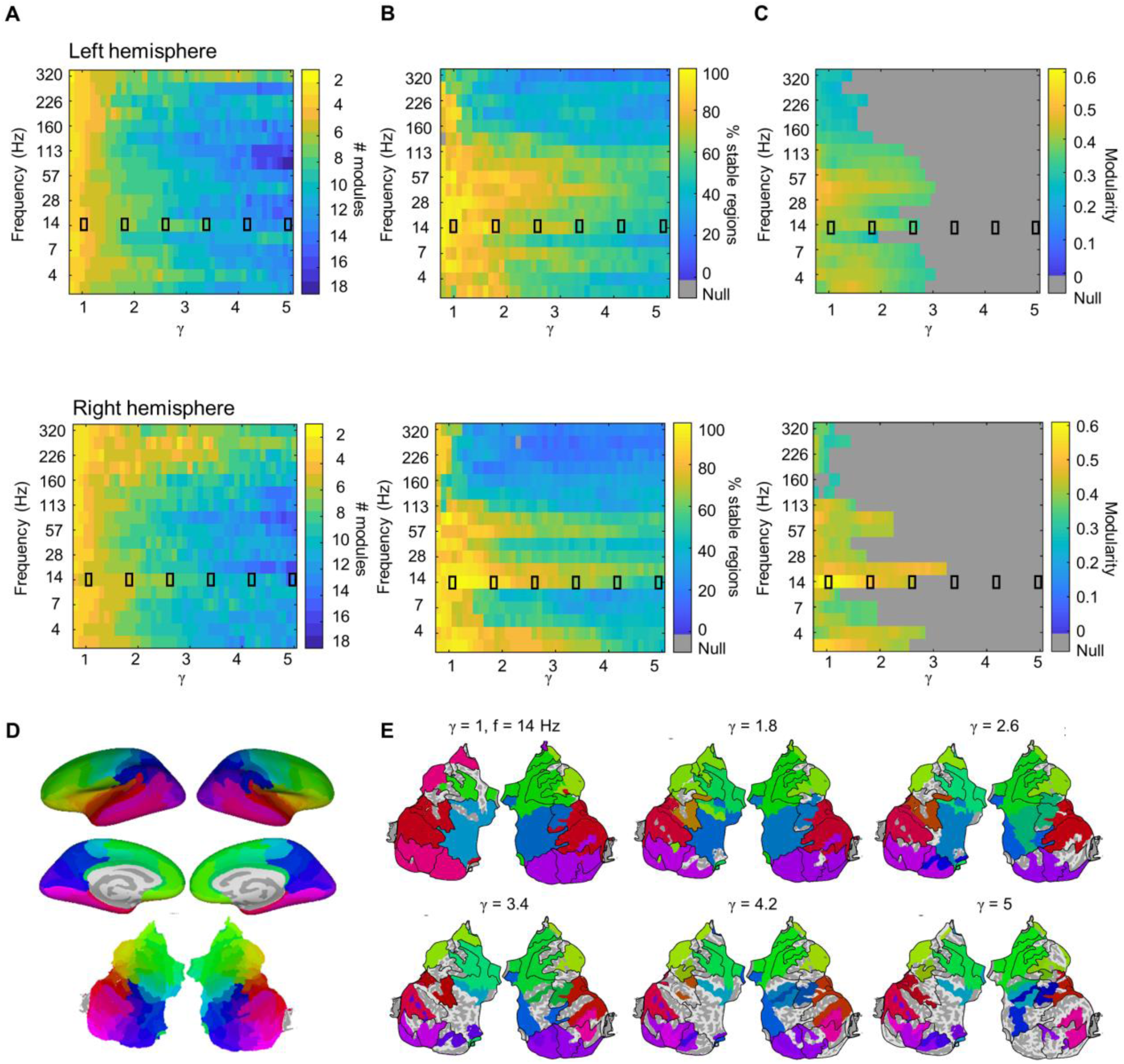
Connectomes of phase-synchronization reveal distinct and stable modules at multiple spatial scales. **A.** Number of identified left and right hemisphere modules, for each combination of spatial scale and filter center frequency. **B.** Percentages of left and right hemisphere regions with stable module assignments, for each combination of spatial scale and filter center frequency. **C.** Modularity for left and right hemisphere, for each combination of spatial scale and filter center frequency. Values below statistical significance are gray. **D.** Translation of colours for each brain region from an inflated to (top) flattened cortical surface (bottom). We performed the transformation from the inflated to flattened cortical surface using the tksurfer FreeSurfer command (Fischl et al. (1999)). **E.** Colour-coded modules for right hemisphere at 14 Hz on flattened cortical surface, at six spatial scales (γ *=* 1 to 5). We converted from HSV to RGB before plotting the modules. Regions with unstable module assignments are gray. Small black rectangles in panels A-C indicate γ values at which modules are visualised in panel E.

We used bootstrapping (*N* = 100 connectomes resampled with replacement) to assess statistical significance of the percentage of brain regions with stable module assignments, and shuffling (*N* = 100 shuffled connectomes) to assess statistical significance of modularity of the original connectomes. Since the outcome of permutation-based significance tests can be sensitive to the number of samples used, we evaluated the robustness of our results to the number of samples used to assess statistical significance. To do this, we compared *z*-scores of the percentages of stable brain regions we obtained with the original 100 bootstrapped connectomes to the corresponding *z*-scores with 1000 bootstrapped connectomes. Similarly, we compared the z-scores of the modularity we obtained with the original 100 randomized connectomes to the corresponding *z*-scores with 1000 randomized connectomes. In the original analysis, the *z*-scores were converted to *p*-values, from which we assessed statistical significance. We found that the *z-*scores of percentages of stable regions for 100 and 1000 bootstrapped connectomes were highly correlated (0.9 for left hemisphere and 0.96 for right hemisphere). Similarly, the *z*-scores of modularity of the connectomes for 100 and 1000 randomized connectomes were highly correlated (0.99 for both left and right hemispheres). These results demonstrate that the results of our permutation-based tests on statistical significance of the identified modules are robust to the number of samples used.

For a given frequency, we illustrate modules on flattened projections of the cortical surface (Fischl et al. (1999)) (Figure 3D). We assigned colours to modules displayed on the flattened cortical surfaces using the following procedure: 1.) Collapsing the set of region *x*-*y* coordinates to a single hemisphere by first flipping all right-hemispheric coordinates about the *y*-axis and estimating the average of *x*-*y* coordinates of the left hemisphere and (flipped) right hemisphere, for each brain region. 2.) Centering the *x*-*y* coordinates by subtracting the mean *x* and *y* coordinates. 3.) For each brain region, estimating Euclidean distance from the (0,0) center, rescaled between 0.6 and 1, and estimating angle from the (0,0) center using the arctan function, rescaled between 0 and 1. 4.) Assigning the colour of each brain region by the Hue Saturation Luminance (HSV) scheme, setting hue as the rescaled angle, luminance as the rescaled distance, and saturation as 1. 5.) Assigning module colours using the HSV scheme, setting hue as the circular mean of angles of constituent regions, rescaled between 0 and 1, saturation as 1, and luminance as the mean of the rescaled distances from the region centers.

At a representative frequency of 14 Hz, modules comprised superior-frontal, inferior-frontal, temporal, parietal and occipital regions at a coarse spatial scale (γ = 1.8). The module of temporal regions split into modules of superior and inferior-temporal regions at finer spatial scales (γ = 2.6) (Figure 3E).

### 3.3 Modules in connectomes of phase-synchronization cluster into canonical frequency bands

Neuronal activity from brain regions fall into distinct frequency bands, *e.g.* delta (1–4 Hz), theta (4– 8 Hz), alpha (8–12 Hz), beta (12–30 Hz) and gamma (30–80 Hz), each with specific behavioural correlates (Buzsáki & Moset (2013), Zhou et al. (2021), Spitzer & Haegens (2017), Zielinski et al. (2019)). Statistical factor analysis on spectral values of EEG activity from brain regions yielded clusters of frequencies that largely corresponded to these canonical frequency bands (Lopes da Silva (2013)), but a data-driven clustering of modules at different frequencies has not been performed. We determined if the identified modules clustered into statistically distinct sets of frequencies. To do this, we first generated matrices of module similarity, between modules at every pair of frequencies, for left and right hemispheres separately. Then, we applied multi-slice community detection (Mucha et al. (2010)) to identify sets of frequencies for which modules were highly similar, for both left and right hemispheres (Figure 4). These module similarity matrices were weighted averages of matrices of module similarity at individual spatial scales, where the weights were specified by the number of frequencies for which the connectomes had statistically significant modular organisation at that spatial scale. We found multiple statistically significant (*p* < 0.05, FDR-corrected, permutation test, *N* = 100) groupings of between two and thirteen frequency bands. For further analysis, we used the groupings into three frequency bands and six frequency bands, though we note that other equally valid groupings could be used. The statistically significant clustering into three frequency bands (γ_multislice_ = 1.1, ω = 0.2–1) comprised sets of adjacent filter center frequencies, 3–14 Hz, 20–113 Hz and 135–320 Hz (Figure 4, dashed red line boxes). Similarly, the statistically significant clustering into six frequency bands (γ_multislice_ = 1.25, ω = 0.2–1) comprised sets of adjacent filter center frequencies, 3–4 Hz, 5–10 Hz, 14–20 Hz, 28–57 Hz, 80–113 Hz and 135–320 Hz (Figure 4, solid black line boxes). Notably, we found an identical clustering into six frequency bands (Supplementary material, Figure S3, solid black line boxes) when we applied equal (unit) weights to the matrices of module similarity at all spatial scales, and the clustering into three frequency bands was also highly similar (3–10 Hz, 14–80 Hz, 113–320 Hz) (Supplementary material, Figure S3, dashed red line boxes). The clustering into six sets of frequencies yielded frequency bands that are close to canonical frequency bands observed in prior literature, *i.e.*, delta (3–4 Hz), theta/alpha (5–10 Hz), beta (14–20 Hz), low gamma (28–57 Hz), high gamma (80–113 Hz) and high-frequency oscillations (135–320 Hz) respectively (Lopes da Silva (2013), Arnulfo et al. (2020)). Thus, the identified modules cluster into statistically distinct sets of frequencies, which map to canonical frequency bands.

**Figure 4.**
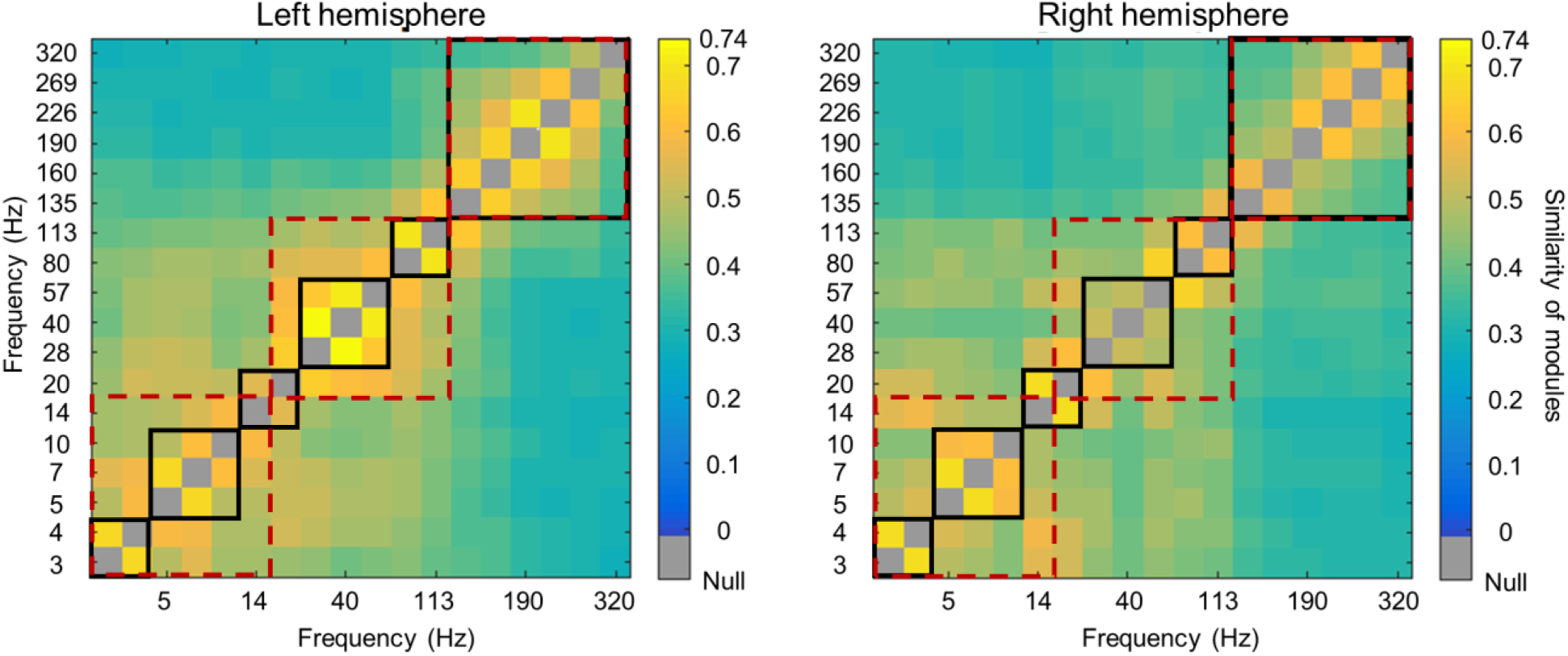
Modules in connectomes of phase-synchronization cluster into canonical frequency bands. Matrices of module similarity, between modules at every pair of frequencies, for left and right hemispheres. Statistically significant clustering common to both hemispheres, into three frequency bands (dashed red outline), *i.e.* 3–14 Hz, 20–113 Hz and 135–320 Hz and into six frequency bands (black outline), *i.e.* 3–4 Hz, 5–10 Hz, 14–20 Hz, 28–57 Hz, 80–113 Hz and 135–320 Hz, are shown.

### 3.4 Modules in connectomes of phase-synchronization comprise anatomically contiguous regions

Module-like structures identified in resting-state fMRI, such as the default mode, fronto-parietal, ventral-and dorsal-attention systems include anatomically non-contiguous regions (Beckmann et al. (2005), van den Heuvel & Pol (2010)). We investigated if modules in connectomes of phase-synchronization similarly comprised anatomically non-contiguous regions for the statistically significant grouping into three and six frequency bands, at different spatial scales (Figure 5). For the grouping into three frequency bands (3–14 Hz, 20–113 Hz and 135–320 Hz), we in fact found the modules comprised anatomically contiguous regions for the 3–14 Hz and 20–113 Hz frequency bands, where the modules respectively comprised frontal, temporal, and parietal regions at a coarse spatial scale (γ = 1). For example, for the 3–14 Hz frequency band, both the left-hemispheric (green) and right-hemispheric (green) modules comprising frontal regions included fronto-marginal gyrus and sulcus, middle frontal gyrus and sulcus, orbital and triangular parts of the inferior frontal gyrus. Similarly, both the left-hemispheric (red) and right-hemispheric (red) modules comprising temporal regions included the temporal pole, inferior temporal gyrus, middle temporal gyrus, superior temporal sulcus and inferior temporal sulcus. Both the left-hemispheric (light blue) and right-hemispheric (dark blue) modules comprising parietal regions included superior parietal gyrus, paracentral gyrus and sulcus, postcentral gyrus and sulcus, and precuneus. At finer spatial scales (γ = 2–4), these modules split into smaller sets of regions, but the brain regions within a module remained anatomically contiguous. For example, the 20–113 Hz frequency band at γ = 2 yielded a left-hemispheric module (brown) including superior temporal regions such as transverse temporal sulcus, anterior transverse temporal gyrus and planum temporale of the superior temporal gyrus, as well as a module (reddish pink) including inferior temporal regions such as the inferior temporal gyrus, inferior temporal sulcus and temporal pole. In contrast to modules for the 3–14 Hz and 20–113 Hz frequency bands however, the modules in the 135–320 Hz frequency band included anatomically non-contiguous regions, across the range of visualised spatial scales (γ = 2–4) (Figure 5) (Arnulfo et al. (2020)). For example, the 135–320 Hz frequency band at γ = 1 yielded a right-hemispheric module (orange) traversing temporal regions such as superior and inferior temporal sulci, parietal regions such as postcentral gyrus and supramarginal gyrus, and occipital regions such as anterior occipital sulcus and middle occipital gyrus. Similar to the modules of the three frequency bands, modules of the six frequency bands (3–4 Hz, 5–10 Hz, 14–20 Hz, 28–57 Hz, 80–113 Hz and 135–320 Hz) comprised anatomically contiguous regions up to 113 Hz, but the modules in the 135–320 Hz frequency band included anatomically non-contiguous regions (Supplementary material, Figures S4–5). Hence, unlike with resting-state fMRI, modules in connectomes of phase-synchronization up to high-gamma frequencies comprised anatomically contiguous regions.

**Figure 5.**
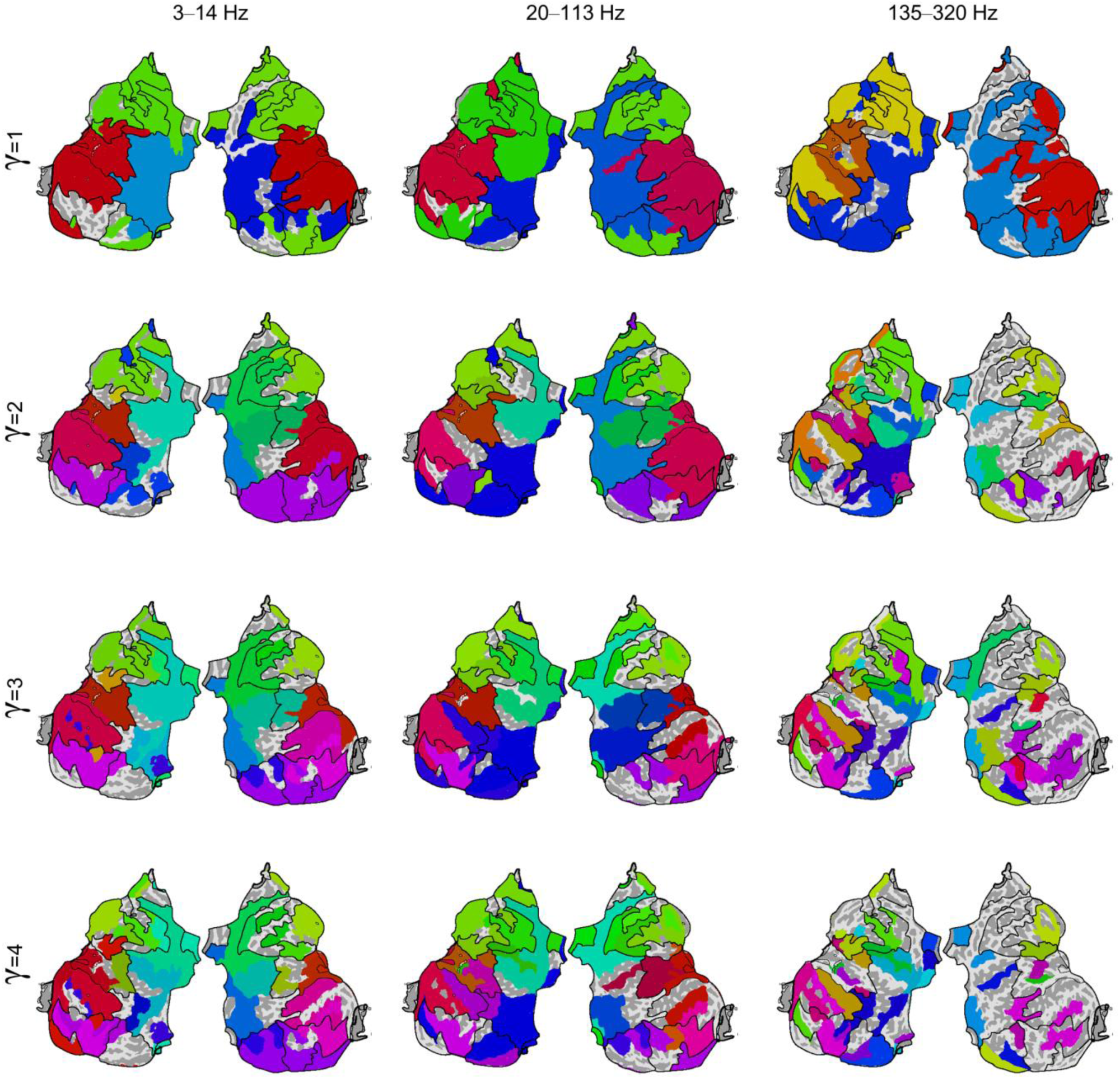
Modules in connectomes of phase-synchronization up to high-gamma frequencies comprise anatomically contiguous regions. Flattened cortical surface representations of modules in connectomes of phase-synchronization for 3–14 Hz, 20–113 Hz and 135–320 Hz, at four spatial scales (γ *=* 1 to 4). Black lines on each flattened surface show outlines of consensus modules, *i.e.* sets of regions assigned to the same module across frequencies and spatial scales.

### 3.5 Modules in connectomes of phase-synchronization comprise functionally related regions

Module-like structures in fMRI functional connectomes, typically recognized as resting-state networks or functional brain systems, comprise regions that are concurrently active in tasks relating to specific sensory, motor, or cognitive domains, such as visual, sensorimotor, attentional, and executive control processing (Smith et al. (2009), Power et al. (2011)). Hence, we investigated if modules in connectomes of phase-synchronization also comprised regions that are concurrently active in tasks relating to specific cognitive domains. For this purpose, we used eight consensus modules that represented sets of regions assigned to the same module across frequencies and spatial scales. In the absence of *a priori* knowledge on the number of consensus modules, we set the number as eight to fall within the range of seven to ten reported for their putative fMRI counterparts (Beckmann et al. (2005), Damoiseaux et al. (2006), Yeo et al. (2011), Power et al. (2011)). The eight consensus modules comprised anatomically contiguous regions and respectively included regions in the superior-frontal (bright green), inferior-frontal (pale green), insula (olive), superior-temporal (brown), inferior-temporal (dark pink), parietal (light blue), lateral-occipital (dark purple), and medial-occipital (light purple) cortical areas (Figure 6A). Module colours reflect anatomical location of their constituent regions (see Section 3.2). The consensus modules predominantly resembled modules at the lower frequencies (14–40 Hz) and intermediate spatial scales (γ = 1.5–2.5) (Supplementary material, Figure S6).

**Figure 6.**
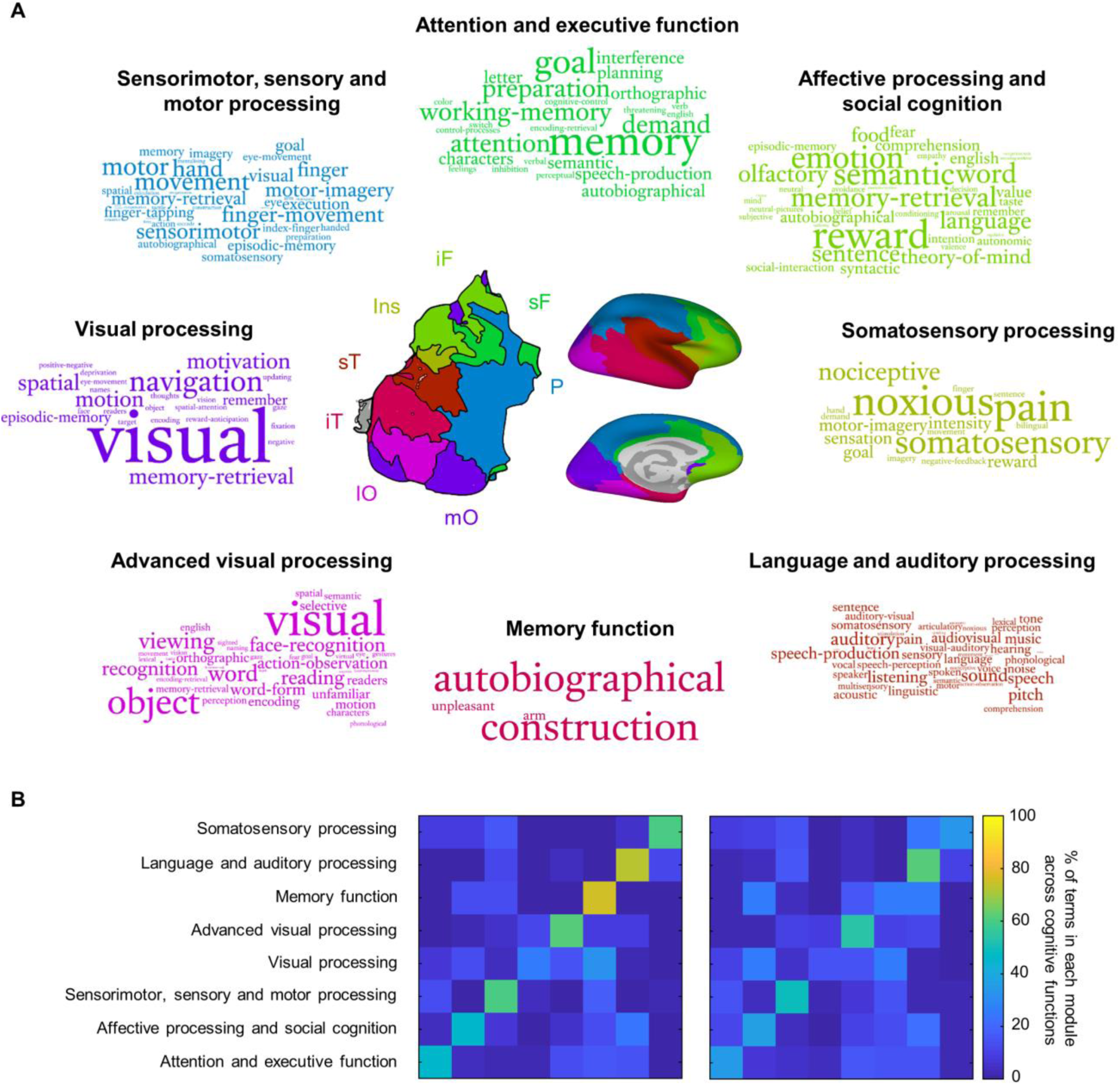
Modules in connectomes of phase-synchronization comprise functionally related regions. **A.** Terms and putative functional roles specific to each of the eight consensus modules displayed in center. Sizes of words are proportional to their frequency of occurrence. sF=superior Frontal, iF=inferior Frontal, Ins=Insula, sT=superior Temporal, iT=inferior Temporal, lO=lateral Occipital, mO=medial Occipital, P=Parietal. **B.** Percentages of terms specific to each module (row) assigned to each of eight cognitive functions (left) and percentages of all terms related to each module (row) assigned to the same cognitive functions (right).

We first used the Neurosynth meta-analyses-based decoding tool (Yarkoni et al. (2011)) to find terms related to perception, cognition and behaviour, selectively associated with each brain region in the Destrieux brain atlas, where we identified each region by its centroid coordinates. These terms were both sensitively and specifically associated to fMRI activation in the corresponding brain regions, according to a large database of fMRI studies. We then identified terms selectively associated with each module by finding terms that occurred more frequently (*p* < 0.05, FDR-corrected, permutation test, *N* = 74) across the regions in a module, compared to equally sized surrogate modules of anatomically contiguous regions. This effectively tested the hypotheses that regions comprising a module serve shared functional roles, even after accounting for their anatomical proximity.

The terms for the superior-frontal module were related to attention and executive function while the inferior-frontal module was associated with affective processing and social cognition (Figure 6A). The parietal module related to sensorimotor, sensory and motor processing, while the medial-occipital and lateral-occipital modules were associated with basic and advanced visual processing respectively. The superior temporal module was related to language and auditory processing, while the inferior temporal module was related to memory function. Finally, the terms for the insula module were associated with somatosensory processing. The results suggest that, similarly to modules in resting-state fMRI, the modules in connectomes of phase-synchronization comprised regions with shared functional roles in task-related processing. The putative functional roles of these modules, inferred from their sets of terms, were in good agreement with overarching functions of their constituent regions (Gazzaniga et al. (2009)).

We sought to further corroborate the functional specificity of modules, *i.e.,* that they are specialised to support particular domains of cognitive functions. To verify this, we determined the percentage of selectively associated terms for each module that could be categorised under every module’s assigned functional role. We compared this against the percentage of all terms for each module, *i.e.,* before FDR-thresholding, that could be categorised under every module’s assigned cognitive function. Functional specificity of modules would be reflected by high percentages of selectively associated terms for each module being assigned to their assigned cognitive function, but the set of all terms for each module being distributed across diverse cognitive functions. As expected, we found high percentages of selectively associated terms for each module were categorised within the cognitive function assigned to them (Figure 6B, left), but the set of all terms for each module were distributed across diverse cognitive functions (Figure 6B, right). These results further verify the functional specificity of the identified modules.

### 3.6 Robustness of results to potential confounds

The modules identified might be influenced by a number of potential confounds, for *e.g.* the community detection method used to identify modules. Hence, we investigated the robustness of the identified modules to several potential confounds. These tests revealed that the modules identified were robust to 1.) the specific sets of SEEG contact-pairs used to generate the group-level connectomes, 2.) the community detection method used to identify modules, 3.) the filter banks used to isolate neuronal activity from different frequencies, 4.) the criterion for the minimum number of SEEG contact-pairs required to estimate a group-level inter-regional connections 5.) the percentile values used to set the connectome threshold, and 6.) regional amplitude differences. Please see Supplementary Text and Figures S7–14 in Supplementary material for further details.

## 4. Discussion

Modules in the fMRI connectome comprise distinct sets of connected regions for sensory, motor and cognitive processing (Valencia et al. (2009), Benjaminsson et al. (2010), Yeo at al. (2011), Power et al. (2011), Lee et al. (2012)). In this study, we investigated whether connectomes of phase-synchronization among meso-and macroscale assemblies of neuronal oscillations exhibit a modular architecture. We used intracerebral SEEG data from 67 subjects to generate connectomes of phase-synchronization (Arnulfo et al. (2020)) which are negligibly affected by volume conduction (Arnulfo et al. (2015a)). We found that connectomes of phase-synchronization exhibited distinct and stable modules at multiple spatial scales at all studied frequencies. Furthermore, data-driven clustering showed that the modules were anatomically similar within canonical frequency bands, *i.e.,* delta (3–4 Hz), theta/alpha (5–10 Hz), beta (14–20 Hz), gamma (28–57 Hz), high-gamma (80–113 Hz) and high frequency (135–320 Hz) bands. In contrast to the modules identified in fMRI, we found that modules up to high-gamma frequency band (80–113 Hz) comprised only anatomically contiguous regions. Importantly, modules comprised brain regions with significantly shared functional roles in *e.g.,* attentional and executive function, language and memory.

### SEEG recordings can be used to identify modules in connectomes of phase-synchronization

Despite the millimeter scale anatomical specificity and high signal-to-noise ratio (SNR) offered by intra-cranial EEG methods like Electrocorticography and SEEG (Parvizi & Kastner (2018)), their sparse spatial coverage and artefacts due to epileptogenic activity have militated against their use to identify modules in connectomes of phase-synchronization. Our results demonstrate the viability of combining SEEG recordings with state-of-the-art methods to identify modules in connectomes of phase-synchronization. We counteracted sparse SEEG coverage by pooling data from 67 subjects and addressed epileptogenic artefacts by removing SEEG contacts and data segments potentially containing epileptic artefactual activity. Further, we used automated procedures to overcome the problem of assigning SEEG contacts to brain regions and used closest-white-matter referencing to minimise volume conduction, to accurately estimate connectomes of phase-synchronization. Finally, we combined consensus clustering with community detection to identify modules in the connectomes despite the presence of missing connections. A recent MEG study (Zhigalov et al. (2017)) used a similar procedure with a smaller cohort (*N* = 27) to estimate the connectome of phase-synchronization, but did not identify modules in these due to the high proportion of missing connections. A recent Electrocorticography (ECoG) study (Kucyi et al. (2018)) measured amplitude correlations between a number of brain regions, but lacked the spatial coverage to estimate the connectome or modules in the connectome. Hence, our study is the first to our knowledge to harness the high SNR and fine anatomical specificity of intra-cranial EEG to study the modular organization of the connectome of phase-synchronization.

It should be mentioned that the yet incomplete coverage offered by SEEG combined with connectome thresholding, might result in missing modules comprising sets of distant (> 90 mm) brain regions. This should be considered when weighing the strengths and limitations of our approach. However, we do reiterate that our investigation of the coverage offered by our method revealed that neither the positions of the SEEG shafts nor thresholding the connectome, precluded identifying modules comprising distant brain regions, including brain regions in different functional subdivisions, *i.e.* frontal, parietal, temporal, occipital, limbic and insular cortices. Rather, we found a number of supra-threshold connections between regions in different functional subdivisions. In fact, modules identified at a coarse spatial scale for the 135–320 Hz frequency group, comprised distant brain regions encompassing parietal, temporal and occipital cortices.

Since SEEG measures LFPs, it is limited in its anatomical specificity, in for *e.g.,* reconstructing the detailed microscopic arrangement of transmembrane currents (Einevoll et al. (2013)). However, SEEG’s anatomical specificity at the level of neuronal populations, together with our closest white-matter referencing scheme, enable accurately estimating inter-regional phase-synchronization and identifying modules in connectomes of phase-synchronization. Compared to MEG, SEEG provides higher spatial resolution due to the minimal influence of volume conduction on estimates of phase-synchronization (Arnulfo et al. (2015a)). Further, SEEG does not have different sensitivities to gyral and sulcal sources, and source orientations, but MEG does (Baillet (2017)).

### SEEG reveals novel modules in connectomes of phase-synchronization

Some of the distinct modules we identified with SEEG have not previously been observed with either fMRI or MEG. The relationship between fMRI connectivity to electrophysiology is multi-factorial, including contributions from both amplitude correlations and phase-synchronization, in multiple frequency bands (Shafiei et al. (2022)). Hence, we do not expect a one-to-one correspondence between the modules we identified in SEEG connectomes of phase-synchronization, and the modules reported with fMRI. We identified modules comprising superior frontal regions, inferior frontal regions, superior temporal regions, inferior temporal regions, parietal regions, insula, lateral occipital regions and medial occipital regions. Modules comprising occipital regions and temporal regions have been identified in resting-state fMRI (Benjaminsson et al. (2010), Yeo et al. (2011), Power et al. (2011)). However, we identified separate modules of medial occipital and lateral occipital regions compared to a single module of occipital regions reported in fMRI, and separate modules of superior temporal and inferior temporal regions compared to a single module of temporal regions reported in fMRI. Further, we identified separate modules of superior frontal and inferior frontal regions, rather than the module of fronto-parietal regions reported in fMRI. Finally, we identified a module of parietal regions, and a module of regions in the insula, both of which have not been previously reported in fMRI. Each of these SEEG modules comprised anatomically contiguous regions in contrast to, for *e.g.*, attentional or default-mode brain systems identified with fMRI, which include regions distributed across frontal, parietal, and temporal cortices (Benjaminsson et al. (2010), Yeo et al. (2011), Power et al. (2011)). The only partial overlap in modules we identified with SEEG to those reported in fMRI is in agreement with the weak correspondence between fMRI connectomes to electrophysiological connectomes of phase-synchronization estimated from MEG data (Shafiei et al. (2022)). Correspondence between fMRI and electrophysiological connectomes was highest in sensory and motor cortices rather than associative cortex (Shafiei et al. (2022)), much the same as the modules we identified with SEEG comprising sensory or motor regions, *e.g.*, the module of superior temporal regions (auditory), corresponding to the module of temporal regions in fMRI data (auditory), but there being no such correspondence for modules comprising associative brain regions.

Previous MEG studies have identified module-like structures representing sets of brain regions whose oscillation amplitude envelopes in specific frequency bands are correlated (Brookes et al. (2011), de Pasquale et al. (2010)). Notably, these studies have demonstrated a strong correspondence to modules identified in fMRI, such as a module of fronto-parietal regions (fronto-parietal control), a module of occipital brain regions (visual) and a module comprising regions in the default mode brain system (Brookes et al. (2011)). Correlation between oscillation amplitude envelopes of brain regions is known to be physiologically distinct to synchronization between oscillation phases (Engel et al. (2013)), and to also exhibit different patterns of inter-regional connectivity (Siems & Siegel (2020)). Hence, we did not expect a strong correspondence between the modules we identified, and previously reported module-like structures of regions whose oscillation amplitude envelopes were correlated. We observed a partial correspondence for a single module - while a previous study (Brookes et al. (2011)) reported a module comprising occipital regions, we reported separated modules for medial occipital regions and lateral occipital regions. However, we also reported modules comprising superior frontal regions, inferior frontal regions, superior temporal regions, inferior temporal regions, parietal regions and regions in the insula, which have not been previously reported in MEG studies identifying sets of regions whose oscillation amplitude envelopes are correlated.

Results from two MEG studies (Zhigalov et al. (2017), Vidaurre et al. (2018)) investigating module-like structures in connectomes of phase-synchronization, corroborate our identification of modules comprising anatomically contiguous regions up to high-gamma frequencies. Zhigalov et al. (2017) reported distinct modules comprising occipital regions, sensorimotor regions and frontal regions. Another recent MEG study (Vidaurre et al. (2018)) used Hidden-Markov modelling to identify spatially localised “functional states”, including those comprising predominantly occipital regions, sensorimotor regions and frontal regions. The “functional states”, were characterised by short-lived patterns of inter-regional coherence and hence, constituted module-like structures. However, in contrast to these MEG studies, we identified separate modules of superior frontal regions and inferior frontal regions and separate modules of medial occipital regions and lateral occipital regions, and we identified a module of parietal regions including both sensorimotor and posterior parietal regions while both the MEG studies reported modules of only sensorimotor regions. The low-resolution parcellations used with MEG to avoid field spread, might distort modules identified at finer spatial scales. We also identified modules comprising superior temporal regions, inferior temporal regions and regions in the insula, that have not been reported before. These might be observed due to the sensitivity of interaction measures, *e.g.,* Phase Locking Value, to near-zero-lag phase-synchronization when used with SEEG. MEG field spread or EEG volume conduction produce high amounts of spurious phase-synchronization when measures such as Phase Locking Value are applied to MEG or EEG data. In contrast, the fine anatomical specificity of SEEG allows using measures sensitive to near-zero-lag phase-synchronization, which then reveal novel sets of regions functionally interacting during resting-state.

Evidence from animal electrophysiology (Leopold et al. (2003)) as well as human SEEG recordings (Arnulfo et al. (2015a)) reveal that strength of phase-synchronization decreases with increasing inter-site distance, which is consistent with the presence of modules comprising anatomically contiguous regions. We also observed modules at frequencies higher than 113 Hz to comprise spatially distant regions. These results are consistent with evidence from intra-cranial EEG recordings (Arnulfo et al. (2020), Vaz et al. (2019), Khodagholy et al. (2017)), demonstrating long-distance phase-synchronization at frequencies exceeding 100 Hz. Phase-synchronization from 113–320 Hz is proposed to reflect broadcasting and transmission of information through High Frequency Oscillations (HFOs), which are generated in deep cortical layers (Arnulfo et al. (2020)).

### Modules at multiple spatial scales consistent with hierarchical organization

Our study is the first to report modular organization at multiple spatial scales in connectomes of phase-synchronization. The module of frontal regions identified at a coarse spatial scale splits into modules of superior frontal regions and inferior frontal regions at a finer spatial scale. Similarly, the module of temporal regions identified at a coarse spatial scale splits into modules of superior temporal regions and inferior temporal regions at a finer spatial scale. This recursive occurrence of sub-modules within modules is consistent with hierarchical modular organization, and has been observed in resting-state fMRI (Meunier et al. (2009)) but not with electrophysiological methods. However, a stricter assessment of hierarchical modular organization requires simultaneously identifying modules at multiple spatial scales. Separately identifying modules at multiple spatial scales, as in the current study, make it difficult to rigorously assess hierarchical modular organization due to the very high number of possible permutations when matching modules across spatial scales.

### Functional specificity of identified modules suggests their behavioural relevance

We used information from an independent database of fMRI studies to infer the functional role of each module. Regions in different modules had shared involvement in cognitive functions of attention and executive function, affective processing and social cognition, somatosensory processing, language and auditory processing, memory function, visual processing, advanced visual processing and sensorimotor processing respectively. The demonstrated functional specificity of these modules suggests that they operate as distinct brain systems. In line with proposed frameworks on brain function (Tononi et al. (1994), Tononi et al. (1998), Balduzzi & Tononi (2008), Lord et al. (2017), Shine et al. (2018)) strong connections within modules might support segregated information processing (Chan et al. (2014)), while weak connections between modules might support integrated information processing (Deco et al. (2015), Westphal et al. (2017)).

We speculate that the identified modules impose a functional architecture of the connectome during resting-state, which is reorganized to meet task-related demands for segregation and integration. Recent frameworks propose that cognitive function is implemented by integration between modules present in the baseline period (Cole et al. (2014), Wig (2017)). Some fMRI studies have found evidence to support this, in the form of associations between cognitive performance and task-related functional reorganization of the brain to facilitate interaction between modules operating at baseline (Spadone et al. (2015), Shine et al. (2016), Cohen & D’Esposito (2016)). While many MEG/EEG studies have found task-related phase-synchronization in for *e.g.,* studies of attention (Lobier et al. (2018)), somatosensory processing (Hirvonen et al. (2018)) and working memory (Kitzbichler et al. (2011)), there are no studies investigating task-related phase-synchronization as reorganization of the functional architecture imposed by modules during resting-state. Future studies could describe task-related phase-synchronization with reference to the natural framework provided by the identified modules in connectomes of phase-synchronization during resting-state, and related frameworks rooted in electrophysiology have been recently proposed (Sadaghiani et al. (2022)).

Since the modules we identified were in resting-state, we emphasise that they naturally accommodate studies on task-related modulations of phase-synchronization, including those in which the distance between interacting regions is inversely related to the frequency of interaction. For example, Womelsdorf et al. (2006) reported task-related gamma-band of phase-synchronization between macaque visual areas, Salazar et al. (2012) reported task-related long-distance beta-band synchronization between macaque frontal and parietal regions and Gross et al. (2004) reported task-related long-distance beta-band synchronization between human frontal, parietal and temporal brain regions. As per the framework imposed by the modules we identified, the task-related short-distance gamma-band synchronization (Womelsdorf et al. (2006)) might reflected segregated information processing via intra-modular connections while the studies reporting task-related long-distance beta-band synchronization (Salazar et al. (2012), Gross et al. (2004)) might reflect integrated information processing via inter-modular connections. However, the framework also accommodates divergences from the principle of distance between brain regions being inversely related to the frequency of interaction. For example, Buschman et al. (2012) reported task-related short-distance alpha-band and beta-band synchronization between electrodes in macaque dorsolateral prefrontal cortex, Michalareas et al. (2016) reported task-related short-distance alpha/beta-band synchronization between visual areas in human MEG, and Melloni et al. (2007) reported task-related long-distance gamma-band synchronization in human EEG. In these cases, the task-related short-distance alpha/beta-band synchronization (Buschman et al. (2012), Michalareas et al. (2016)) might reflect segregated information processing via intra-modular connections while the task-related long-distance gamma-band synchronization (Melloni et al. (2007)) might reflect integrated information processing via inter-modular connections. Thus, the modules provide a natural framework to interpret results of studies on task-related phase-synchronization.

### Directions for future work

We propose two particularly promising directions to build on this work. While we studied the anatomical composition of each of the modules, we did not investigate the relationships between modules. Studying the balance between intra-modular and inter-modular connections of brain regions within each of the modules might provide clues to the role of the module within the whole-brain system (Guimerà & Amaral (2005)). For example, some modules might serve as “processing systems” while others might play the role of “control systems” (Power et al. (2011)). Another promising direction is to consider other means by which functional segregation might be implemented, in addition to segregation made possible by the modular structure defined by the connection strengths (Dotson et al. (2014)). In particular, the phase-lags of the synchronization between every pair of regions could be studied to determine if for *e.g.,* phase-lags between regions within a module are lower than phase-lags between regions in different modules, thus reinforcing the segregation of information processing imposed by the modular organisation.

## 5. Conclusion

In this study, we combined resting-state SEEG recordings with state-of-the-art methods to accurately identify modules in connectomes of phase-synchronization. We found the modules to predominantly comprise anatomically contiguous regions, unlike modules identified in resting-state fMRI. Importantly, each of the modules comprised regions with shared involvement in specific cognitive functions. Hence, these modules might represent distinct brain systems with particular roles in perceptual, cognitive and motor processing.

## Acknowledgments

The authors gratefully acknowledge the support of Human Brain Project (604102), Sigrid Juselius Foundation and Academy of Finland (J.M.P. project numbers: 253130, 256472, 281414, 296304, 266745. S.P. project numbers: 266402, 266745, 303933, 325404) to complete this project. Further, the authors are grateful to Jonni Hirvonen and Santeri Rouhinen, for help with data processing, and to Dr. Franceso Cardinale and Annalisa Rubino for facilitating the SEEG recordings.

