## Supplementary text and figures for "Modules in connectomes of phase-synchronization comprise anatomically contiguous, functionally related regions"

### 1. Relationship between connectome coverage and connection distance

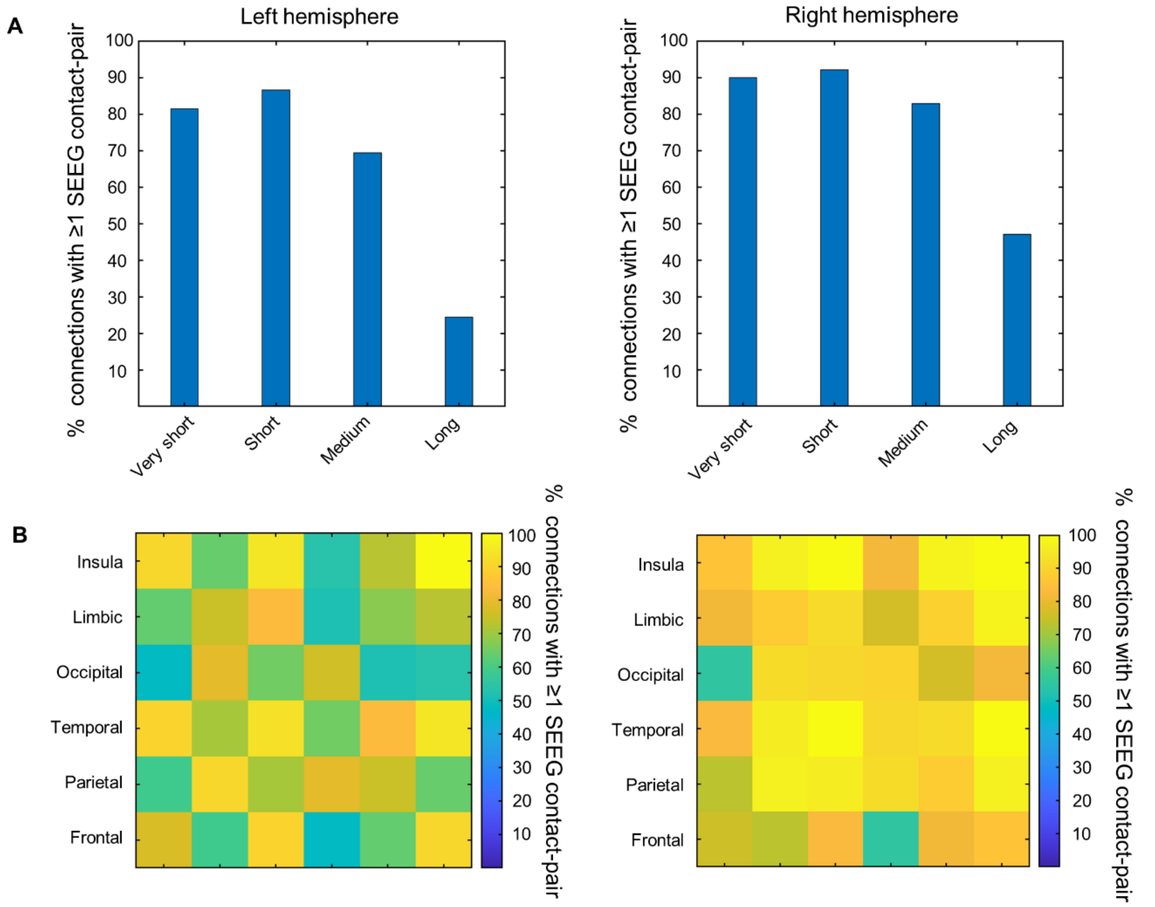


**Figure S1. SEEG recordings provide coverage of inter-regional connections both between proximal brain regions and between distant brain regions A.** Percentage of inter-regional connections sampled by at least one SEEG contact-pair, for very short (< 30 mm), short (30–60 mm), medium (60–90 mm) and long (> 90 mm) distances. **B.** Percentage of inter-regional connections by at least one SEEG contact-pair between every pairwise combination of six functional subdivisions: frontal, parietal, temporal, occipital, limbic and insula.

### 2. Relationship between percentage of supra-threshold connections and connection distance

We identified modules in thresholded connectomes wherein we retained the strengths of all connections in the top 20 percentile while setting others to 0. To check the sampling statistics, we investigated the coverage of just the supra-threshold connections with respect to distance, within each hemisphere. The presence of supra-threshold connections both between proximal and between distant brain regions would allow the community detection method to identify modules comprising both proximal and distant regions, while the presence of supra-threshold connections between only proximal brain regions would limit the community detection method to identifying modules comprising only proximal brain regions. Based on previously reported results, we expected that supra-threshold connections would be more common between proximal brain regions (Leopold et al. (2003), Arnulfo et al. (2015a)), but we also expected a small percentage of supra-threshold connections between distant brain regions. We determined the coverage of supra-threshold connections, at each of the 18 frequencies studied, for both hemispheres, by 1.) estimating the number of supra-threshold connections as a percentage of the number of inter-regional connections sampled by the SEEG contacts, for each of four distances: very short (< 30 mm), short (30–60 mm), medium (60–90 mm) and long (> 90 mm), and 2.) estimating the number of supra-threshold connections as a percentage of the number of inter-regional connections sampled by the SEEG contacts, between regions in every pairwise combination of the following functional subdivisions: frontal, parietal, temporal, occipital, limbic and insular cortices. We found 43 % ± 10 % (mean ± SD) percentage of supra-threshold connections of very-short connections for left hemisphere and 44 % ± 12 % for right hemisphere, while we only found 7 % ± 5 % long-distance supra-threshold connections for left hemisphere and 9 % ± 6 % for right hemisphere (Figure S2A). The higher frequencies had greater percentages of long-distance supra-threshold connections. While the presence of long-distance supra-threshold connections was low, we found several pairs of functional subdivisions, *i.e.* frontal, parietal, temporal, occipital, limbic and insular cortices, between whose regions ~20 % of supra-threshold connections were present (Figure S2B). For example, inter-regional connections between frontal and parietal regions at 14 Hz in the left hemisphere, had 21 % supra-threshold connections. When retaining top 20 percentile strongest connections, 20 % supra-threshold connections are in fact the expected value for the case of no relationship between connection strength and distance. Hence, we find that the thresholding does retain more short-distance than long-distance connections, but that does not preclude identifying modules comprising distant brain regions, including between regions in different functional subdivisions, *i.e.* between regions in frontal, parietal, temporal, occipital, limbic and insular cortices.

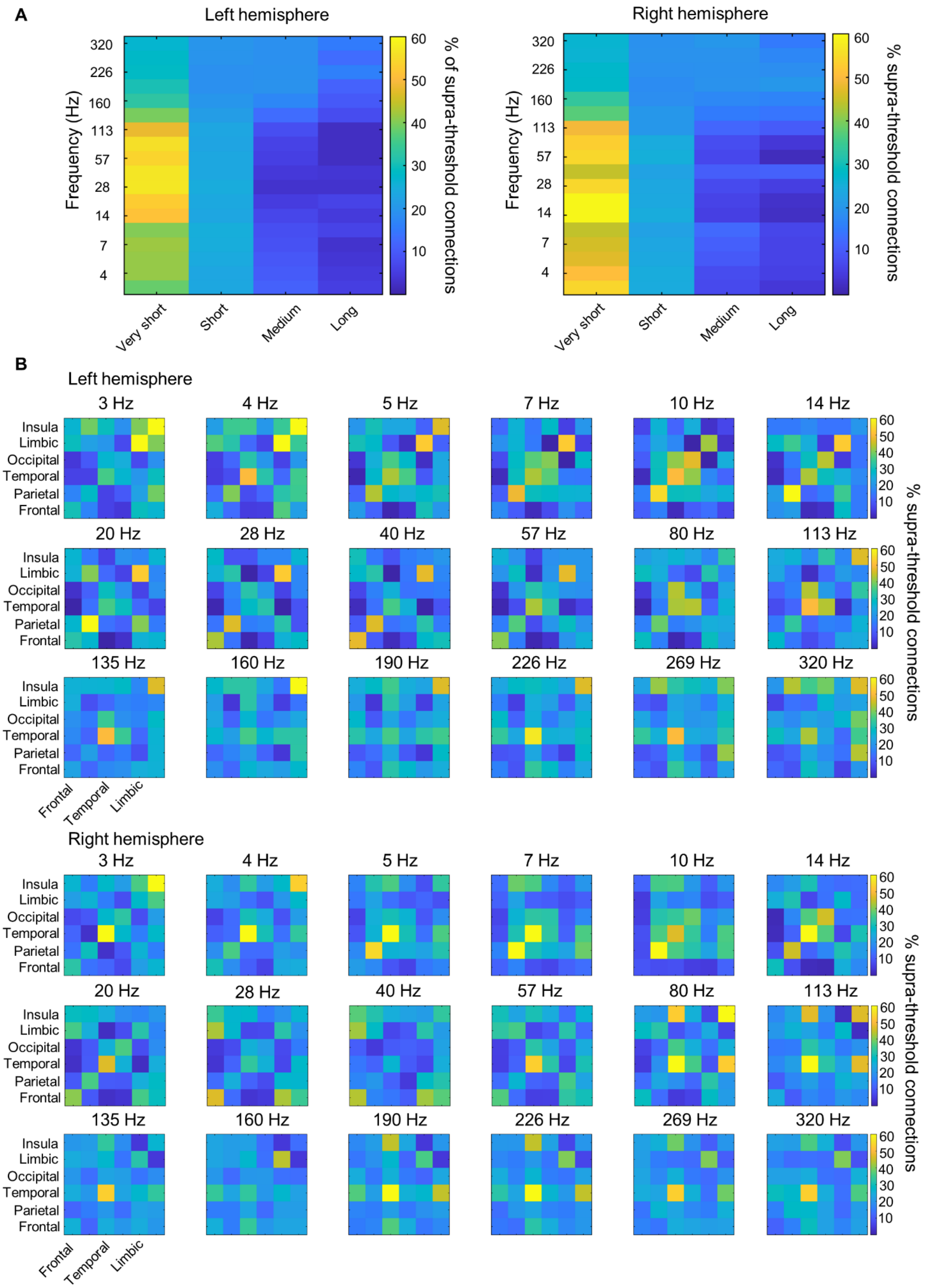
A
Left hemisphere
Right hemisphere
Frequency (Hz)
320
226
160
113
57
28
14
7
4
Very short
Short
Medium
Long
60
50
40
30
20
10
% of supra-threshold connections
% supra-threshold connections
B
Left hemisphere
Right hemisphere
3 Hz
4 Hz
5 Hz
7 Hz
10 Hz
14 Hz
20 Hz
28 Hz
40 Hz
57 Hz
80 Hz
113 Hz
135 Hz
160 Hz
190 Hz
226 Hz
269 Hz
320 Hz
Insula
Limbic
Occipital
Temporal
Parietal
Frontal
Frontal
Temporal
Limbic

**Figure S2. Percentile thresholding retains inter-regional connections both between proximal and between distant brain regions. A.** Percentage of supra-threshold connections, with respect to the total number of sampled connections, at very short (< 30 mm), short (30–60 mm), medium (60–90 mm) and long (> 90 mm) distances, for each of the 18 frequencies studied. **B.** Percentage of supra-threshold connections, with respect to the total number of sampled connections, between regions in every pairwise combination of the six functional subdivisions: frontal, parietal, temporal, occipital, limbic and insula, for each of the 18 frequencies studied.

### 3. Robustness of identified groups of frequencies to weights assigned to module similarity matrices at individual spatial scales

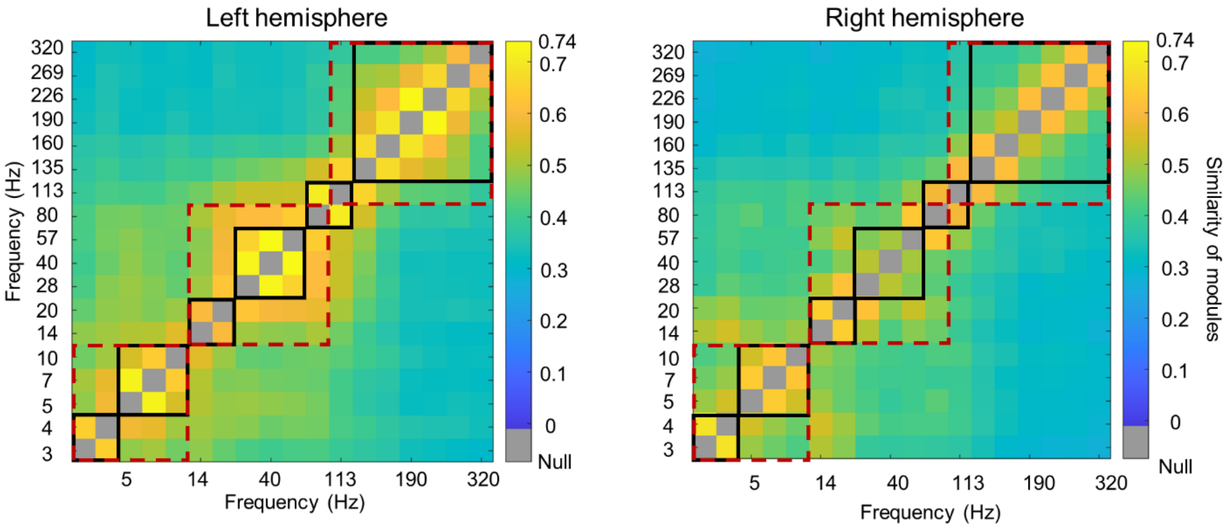


**Figure S3. Clustering of frequencies by module similarity is robust to weighting scheme applied to module similarity at individual spatial scales.** Matrices of module similarity between modules at every pair of frequencies, for left and right hemispheres. Statistically significant clustering common to both hemispheres, into three frequency bands (dashed red outline), *i.e.*, 3–10 Hz, 14–80 Hz and 113–320 Hz and into six frequency bands (black outline), *i.e.*, 3–4 Hz, 5–10 Hz, 14–20 Hz, 28–57 Hz, 80–113 Hz and 135–320 Hz, are shown.

#### 4. Modules for each of the six frequency bands identified by multi-slice community detection

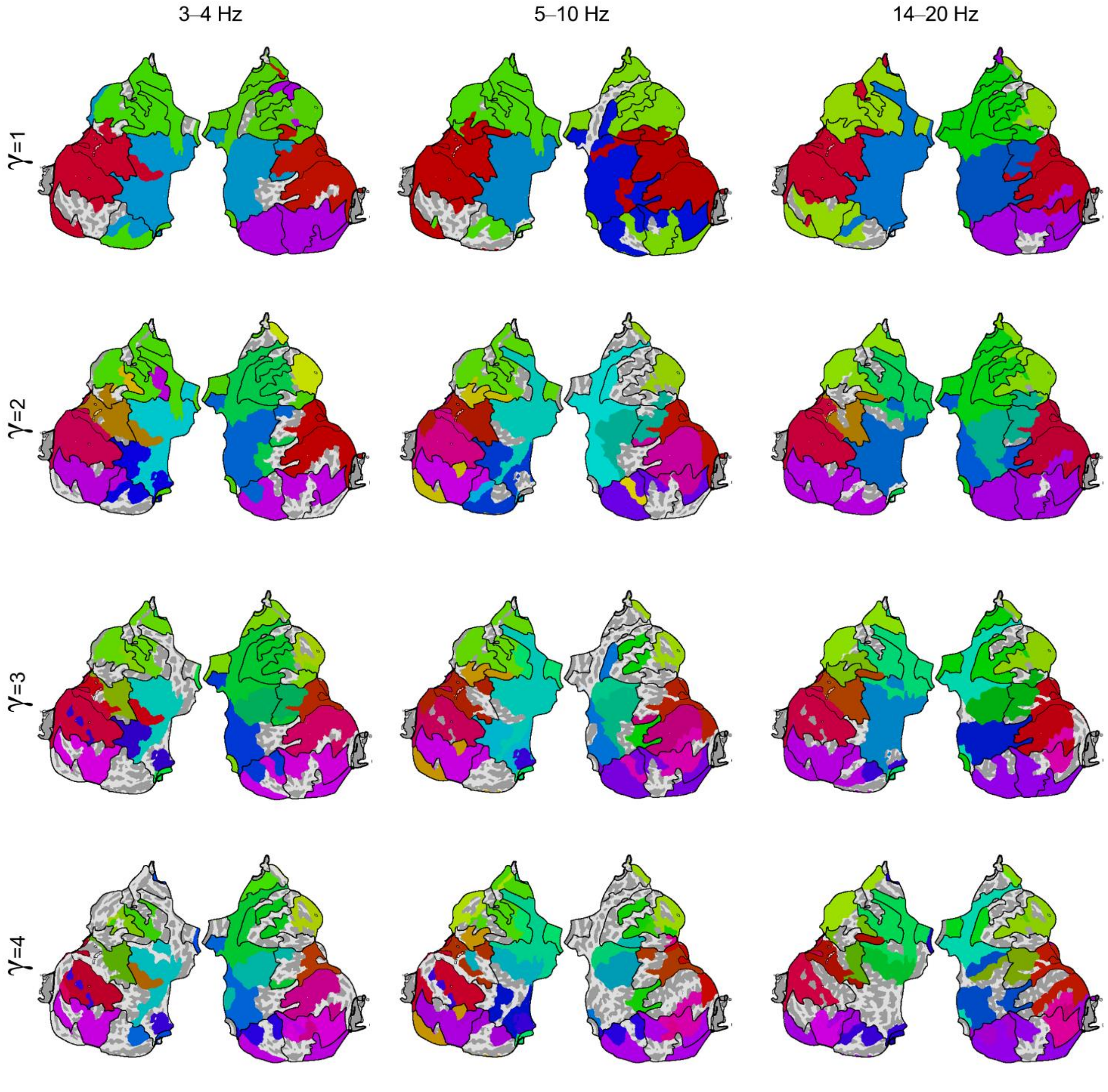


**Figure S4. Modules in connectomes of phase-synchronization for 3–4 Hz, 5–10 Hz and 14–20 Hz comprise anatomically contiguous regions.** Flattened cortical surface representations of modules in connectomes of phase-synchronization for 3–4 Hz, 5–10 Hz and 14–20 Hz, at four spatial scales ($\gamma$ = 1 to 4). Black lines on each flattened surface show outlines of consensus modules, *i.e.*, sets of regions assigned to the same module across frequencies and spatial scales.

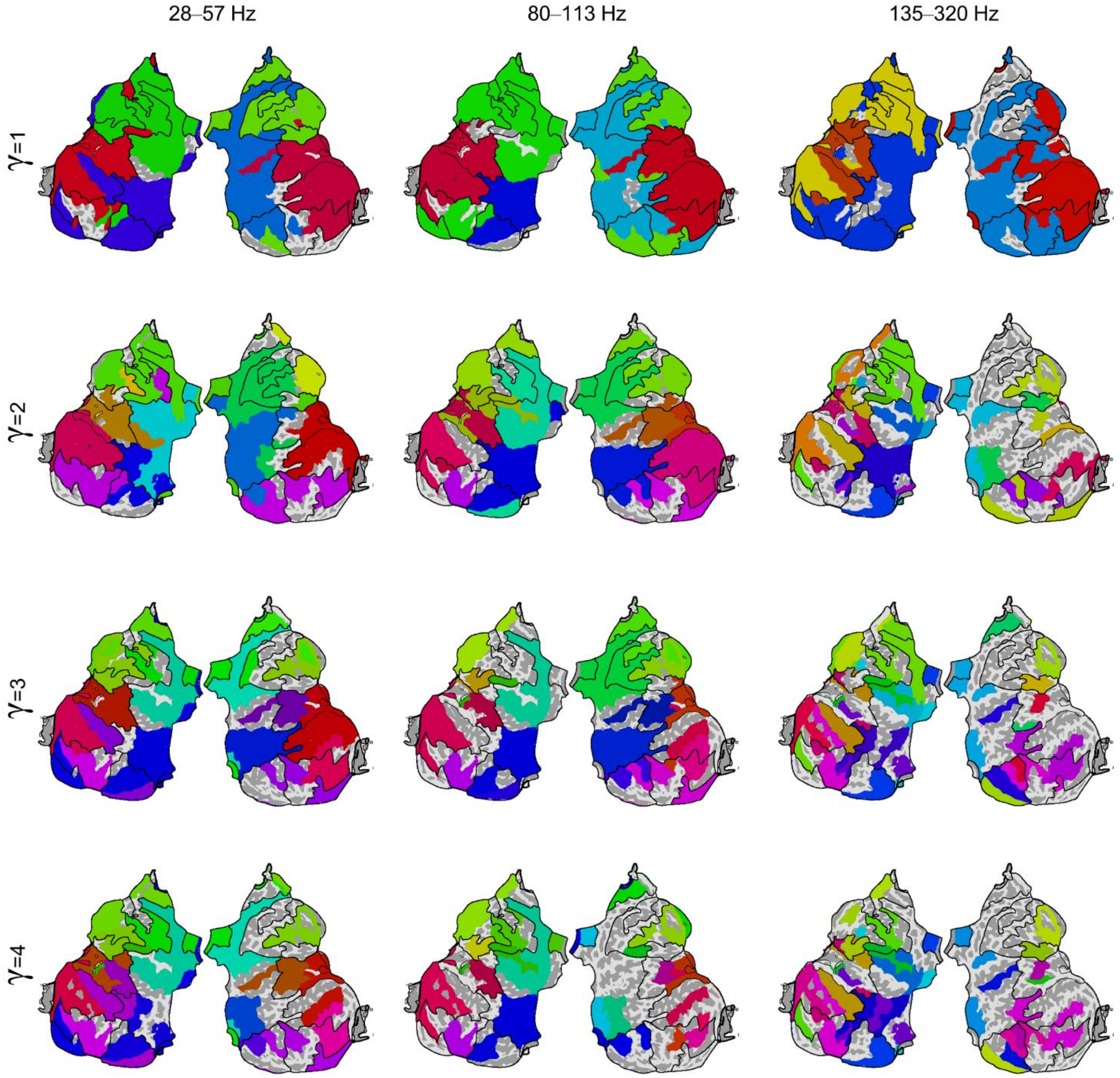


**Figure S5. Modules in connectomes of phase-synchronization for 28–57 Hz, 80–113 Hz but not 135–320 Hz comprise anatomically contiguous regions.** Flattened cortical surface representations of modules in connectomes of phase-synchronization for 28–57 Hz, 80–113 Hz and 135–320 Hz, at four spatial scales ($\gamma$ = 1 to 4). Black lines on each flattened surface show outlines of consensus modules, *i.e.* sets of regions assigned to the same module across frequencies and spatial scales.

#### 5. Similarity between consensus modules and modules at individual frequencies and spatial scales

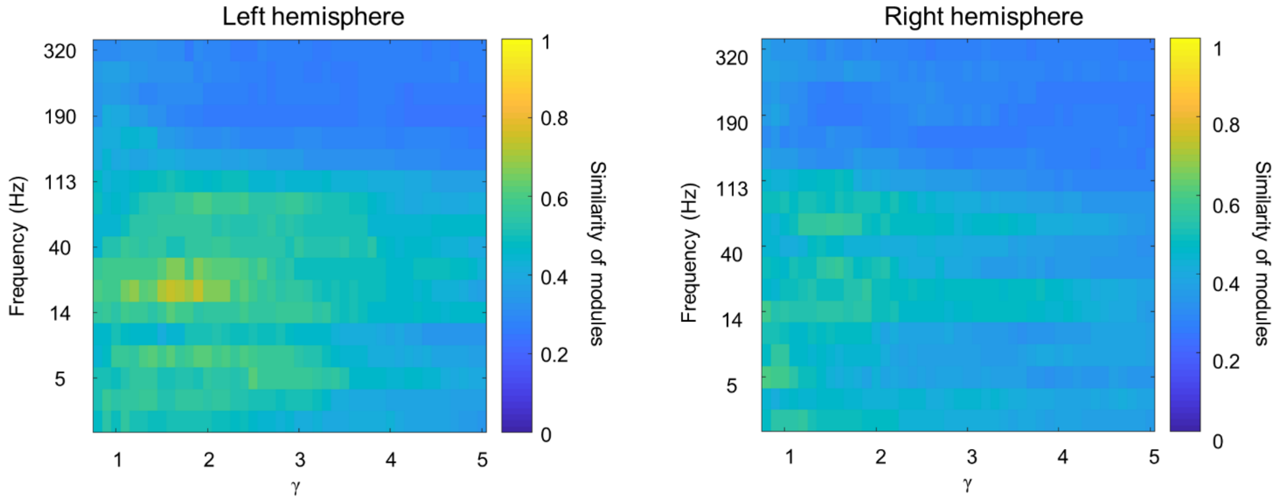


**Figure S6. Consensus modules resemble modules at lower frequencies and intermediate spatial scales.** Similarity between consensus modules and modules at each combination of spatial scale and frequency, for both left and right hemispheres.

#### 6. Robustness of identified modules to potential confounds

We evaluated the robustness of the identified modules to the specific SEEG contact-pairs used to generate the connectomes. To do this, we generated two split connectomes from the original connectome and compared the modules identified from each. Modules identified from the split connectomes were highly similar to each other (Figure S7). Hence, the identified modules were robust to the particular SEEG contact-pairs used to generate the connectomes.

We further evaluated the robustness of the identified modules to the algorithm used for community detection. To do this, we identified modules with Infomap community detection (Rosvall & Bergstrom (2008)) and compared these to the modules we had identified with Louvain community detection. Modules identified by both these methods were highly similar up to high-gamma (113 Hz) (Figure S8). Hence, the identified modules were robust to the algorithm used up to high-gamma frequencies but were algorithm-specific for high-frequency oscillations (135–320 Hz).

The filter banks we used had log-increasing window widths, which offered fine spectral resolution at lower frequencies while simultaneously providing fine temporal resolution of the known-to-be short-lived oscillations at higher frequencies. However, the wide window widths at higher frequencies might allow multiple oscillatory components to be included, potentially confounding the estimation of instantaneous phase which assumes a single oscillatory component. This potential confound would manifest in altering the strengths of estimated phase-synchronization between two brain regions compared to the actual strength of phase-synchronization between these regions. We investigated the robustness of our results to this potential confound by identifying modules on binarized versions of the original connectomes, and comparing these to the modules identified on the original weighted connectomes. Since the potential confound of mis-specified filter banks would manifest in altering the strengths/weights of inter-regional phase-synchronization, highly similar modules identified on the binarized and corresponding weighted connectomes would indicate that our results are robust to this potential confound. We found that modules identified on binarized versions of the connectomes were indeed highly similar to those identified on the original weighted connectomes (Figure S9). Hence, we conclude that our results are robust to the potentially confounding effect of the specified filter banks on the estimation of instantaneous phase.

We imposed no threshold on the minimum number of SEEG contact-pairs required, to estimate an inter-regional connection. We determined the robustness of the modules we identified to this minimum number of SEEG contact-pairs. To do this, we compared the modules we identified on the original connectome, to modules we identified on connectomes generated by imposing a minimum number of 5 SEEG contact-pairs and 10 SEEG contact-pairs. We found modules we identified on the original connectome to be highly similar to modules we identified on connectomes generated by imposing a minimum of 5 SEEG contact-pairs and 10 SEEG contact-pairs (Figure S10–11). Hence, we conclude that the modules we identified are robust to the threshold of the minimum number of SEEG contact-pairs we imposed, to estimate an inter-regional connection.

We thresholded the connectomes before identifying modules by retaining the phase-synchronization strengths of connections in the top 20 percentile, while setting all other connections to 0. We determined the robustness of our results to this choice of this percentile threshold. To do this, we compared the modules we identified on the original connectome after retaining the top 20 percentile connections, to modules identified on connectomes wherein we retained the top 10 percentile and top 30 percentile connections respectively. We found modules identified on the original connectome to be highly similar to modules we identified on connectomes generated by retaining the top 10 percentile and 30 percentile connections (Figure S12–13). Hence, we conclude that the modules we identified are robust to the specific percentile value used to the threshold the connectomes, before identifying modules.

Finally, we investigated if identifying the modules is confounded by the amplitudes of oscillations of individual nodes of the network. We compared modules identified from 67 subject-level networks of phase-synchronization, across all 18 frequencies, at six spatial scales ($\gamma$ = 1, 1.8, 2.6, 3.4, 4.2 and 5), before and after removing amplitude-related differences in strengths of functional connections. The identified modules were highly similar before and after correcting for amplitude-related differences, across all subjects and frequencies, for each of the six spatial scales (mean = 0.92, SD = 0.1) (Figure S14). Hence, the identified modules are not confounded by oscillation amplitudes of individual network nodes.

#### 6.1 Robustness of modules to samples used to generate group-level connectomes

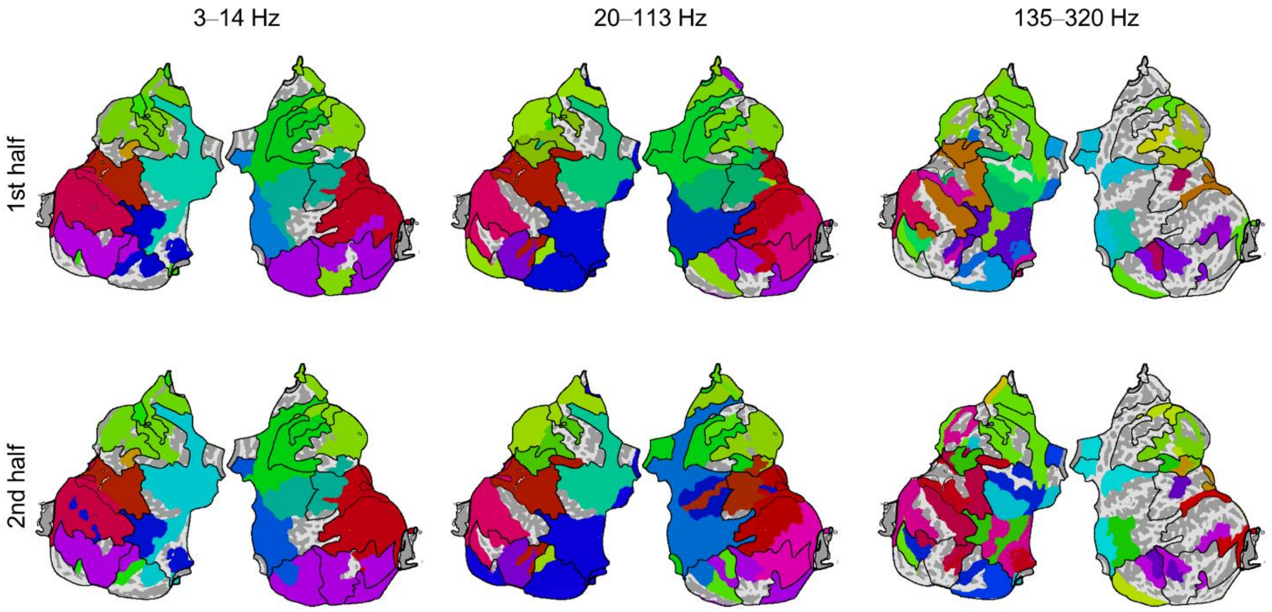


**Figure S7. Modules in split connectomes of phase-synchronization are highly similar.** Flattened cortical surface representations of modules in two split connectomes of phase-synchronization (top and bottom rows) for 3–14 Hz, 20–113 Hz and 135–320 Hz, at a single spatial scale ($\gamma$ = 2). Black lines on each flattened surface show outlines of consensus modules, *i.e.* sets of regions assigned to the same module across frequencies and spatial scales.

#### 6.2 Robustness of modules to community detection method used

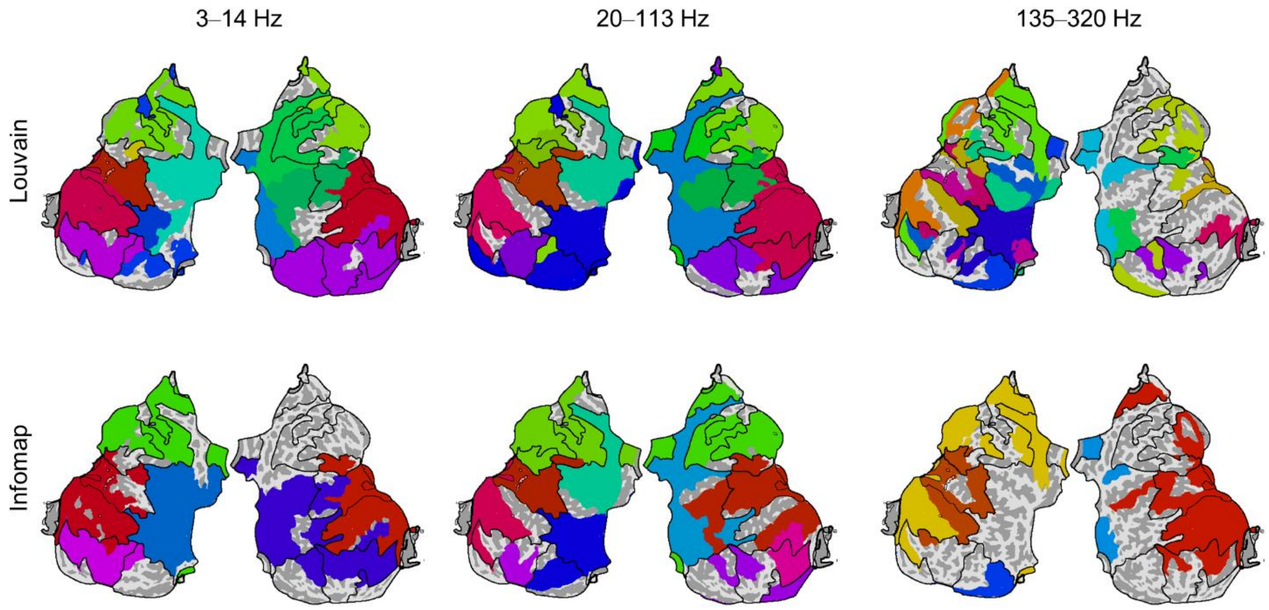


**Figure S8. Modules in connectomes of phase-synchronization similar for Louvain and Infomap community detection, up to high gamma frequency band.** Flattened cortical surface representations of modules in connectomes of phase-synchronization estimated with Louvain (top row) and Infomap (bottom row) community detection, for 3–14 Hz, 20–113 Hz and 135–320 Hz, at a single spatial scale ($\gamma = 2$). Black lines on each flattened surface show outlines of consensus modules, *i.e.* sets of regions assigned to the same module across frequencies and spatial scales.

#### 6.3 Robustness of modules to whether connectome is binary or weighted

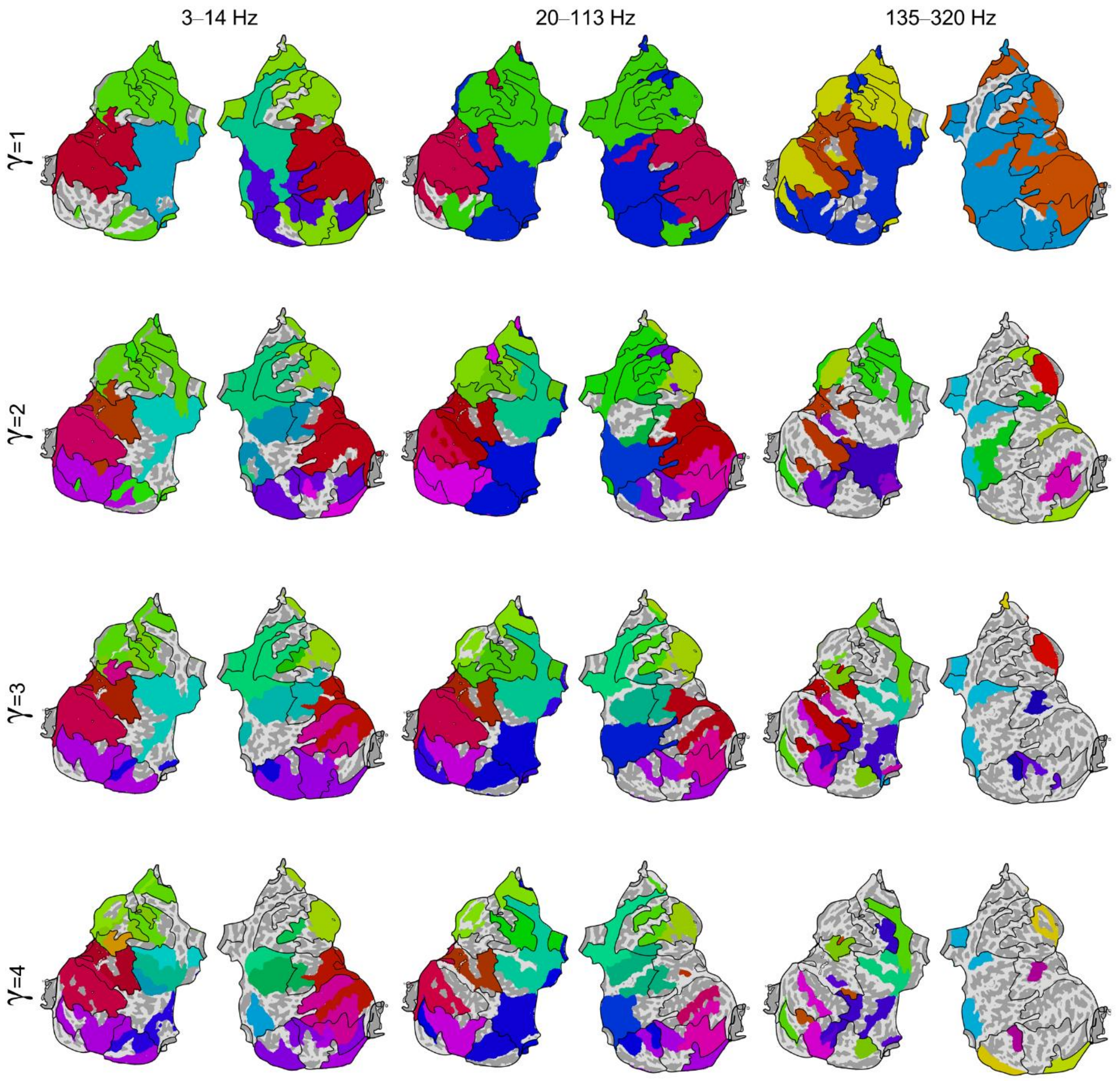


**Figure S9. Modules in binarized connectomes of phase-synchronization are similar to modules in original weighted connectomes of phase-synchronization.** Flattened cortical surface representations of modules in binarized connectomes of phase-synchronization for 3–14 Hz, 20–113 Hz and 135–320 Hz, at four spatial scales (γ = 1 to 4). Black lines on each flattened surface show outlines of consensus modules, *i.e.* sets of regions assigned to the same module across frequencies and spatial scales.

#### 6.4 Robustness of modules to minimum number of samples required to estimate group-level connection

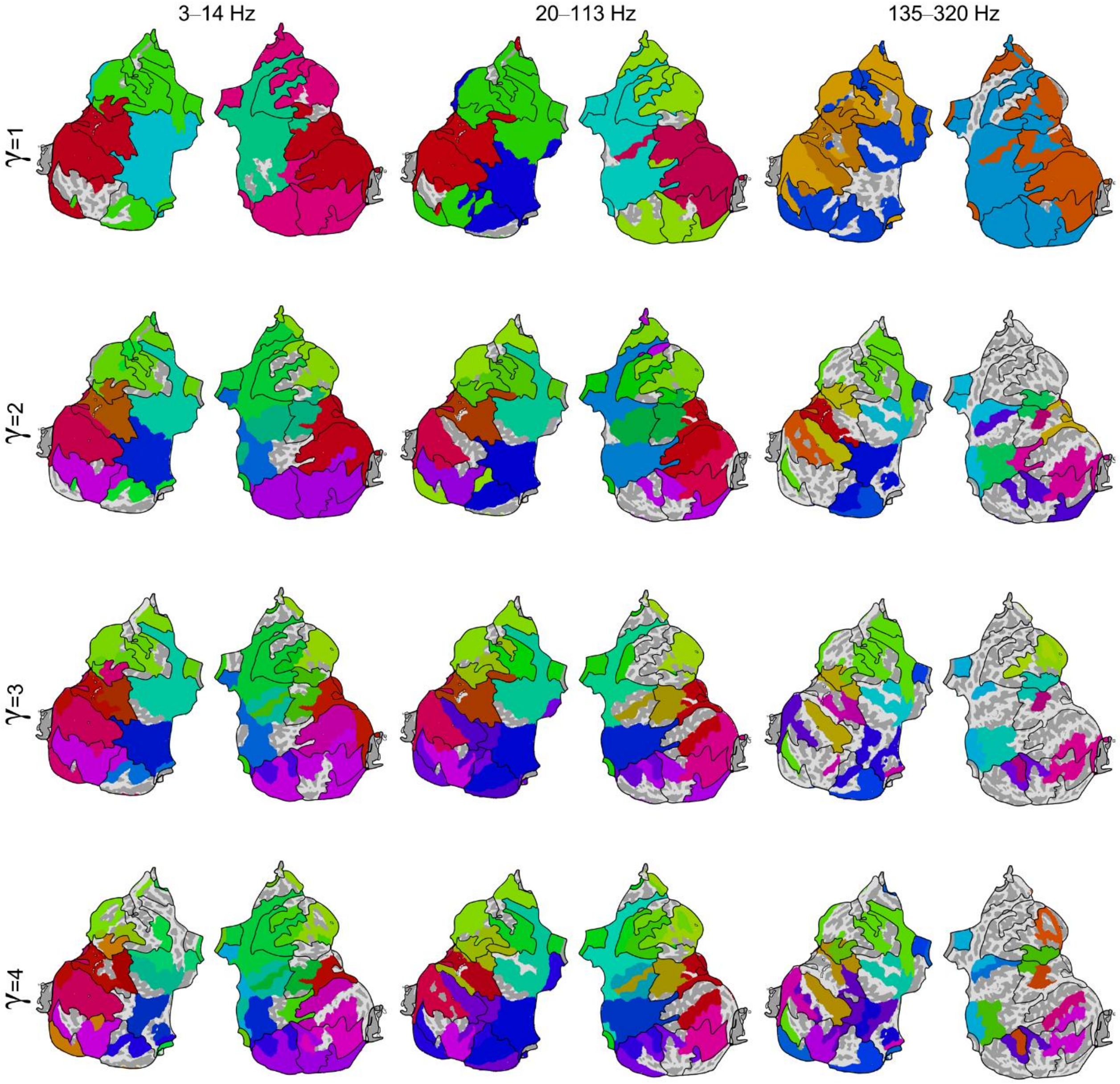


**Figure S10. Modules in connectomes of phase-synchronization generated with criterion on minimum of 5 samples per connection, similar to modules in original connectomes of phase-synchronization.** Flattened cortical surface representations of modules in connectomes of phase-synchronization for 3–14 Hz, 20–113 Hz and 135–320 Hz, at four spatial scales ($\gamma$ = 1 to 4). Each inter-regional connection was subject to the criterion of a minimum of 5 SEEG contact-pairs traversing that pair of brain regions. Black lines on each flattened surface show outlines of consensus modules, *i.e.* sets of regions assigned to the same module across frequencies and spatial scales.

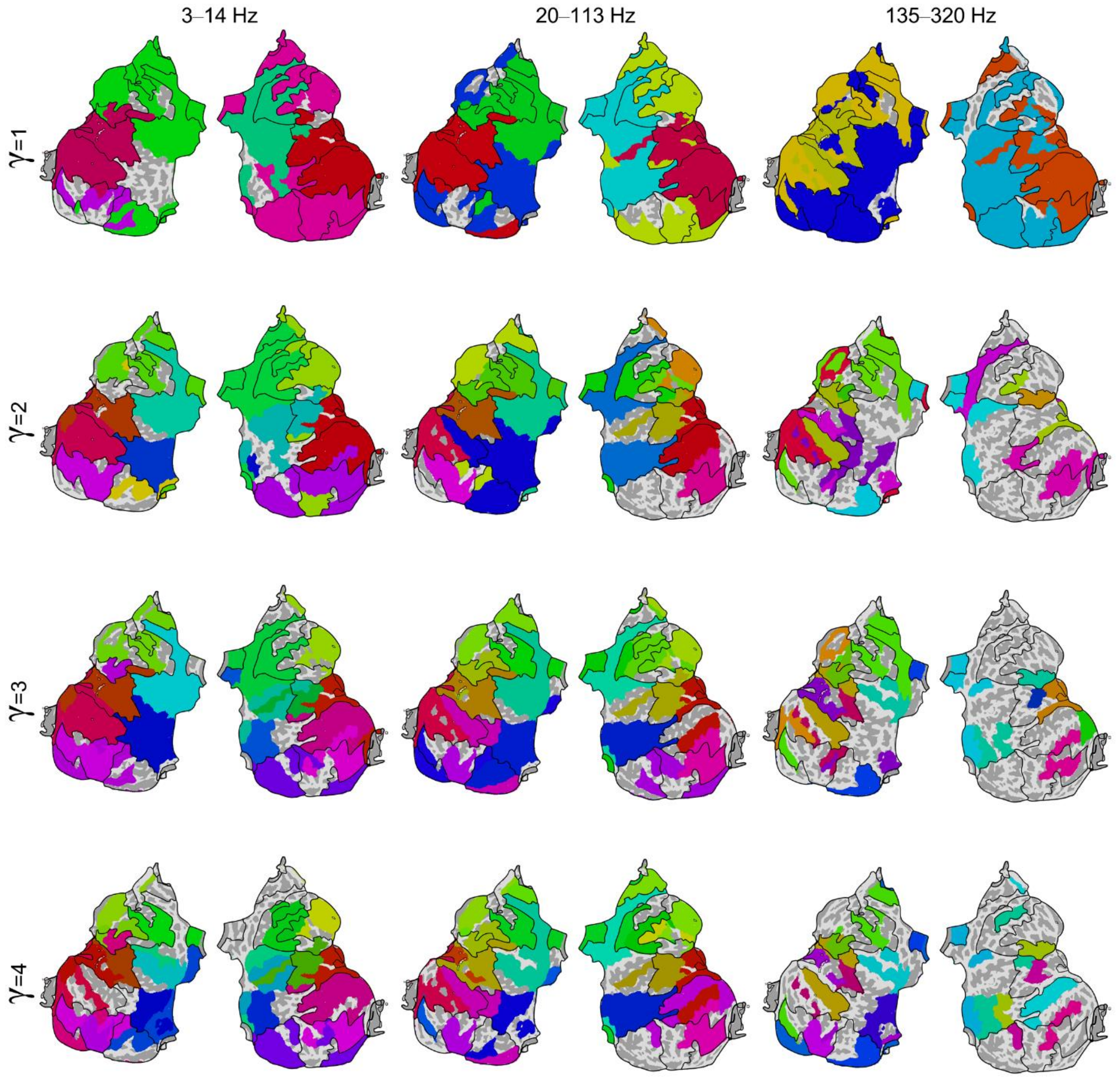


**Figure S11. Modules in connectomes of phase-synchronization generated with criterion on minimum of 10 samples per connection, similar to modules in original connectomes of phase-synchronization.** Flattened cortical surface representations of modules in connectomes of phase-synchronization for 3–14 Hz, 20–113 Hz and 135–320 Hz, at four spatial scales (γ = 1 to 4). Each inter-regional connection was subject to the criterion of a minimum of 10 SEEG contact-pairs traversing that pair of brain regions. Black lines on each flattened surface show outlines of consensus modules, *i.e.* sets of regions assigned to the same module across frequencies and spatial scales.

#### 6.5 Robustness of modules to percentile value for connectome thresholding

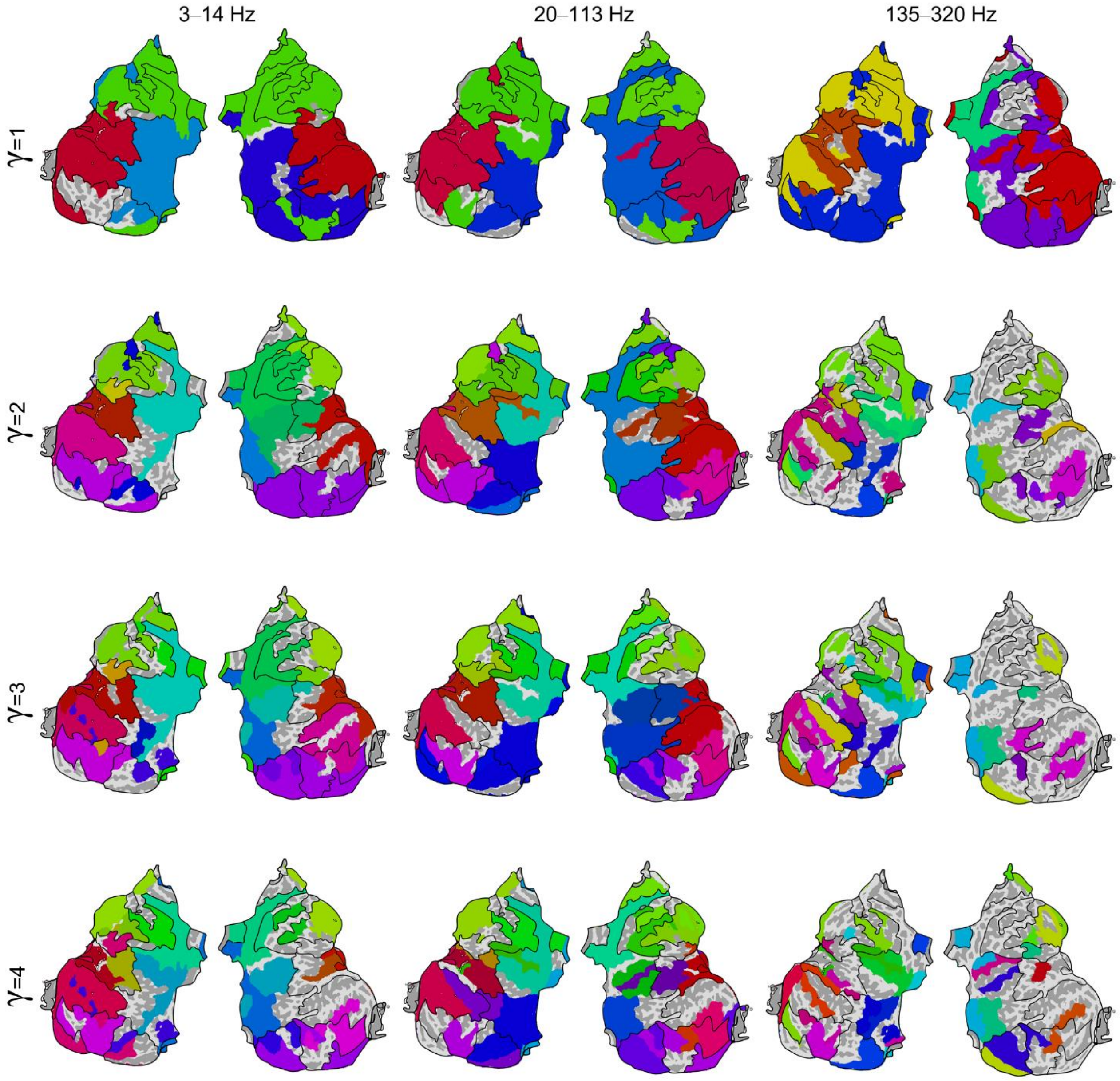


**Figure S12. Modules in connectomes of phase-synchronization generated by retaining top 30 percentile of connections, similar to modules in original connectomes of phase-synchronization.** Flattened cortical surface representations of modules in connectomes of phase-synchronization for 3–14 Hz, 20–113 Hz and 135–320 Hz, at four spatial scales ($\gamma$ = 1 to 4). Connectomes were generated by retaining weights of top 30 percentile of inter-regional connections, while setting all other connections to 0. Black lines on each flattened surface show outlines of consensus modules, *i.e.* sets of regions assigned to the same module across frequencies and spatial scales.

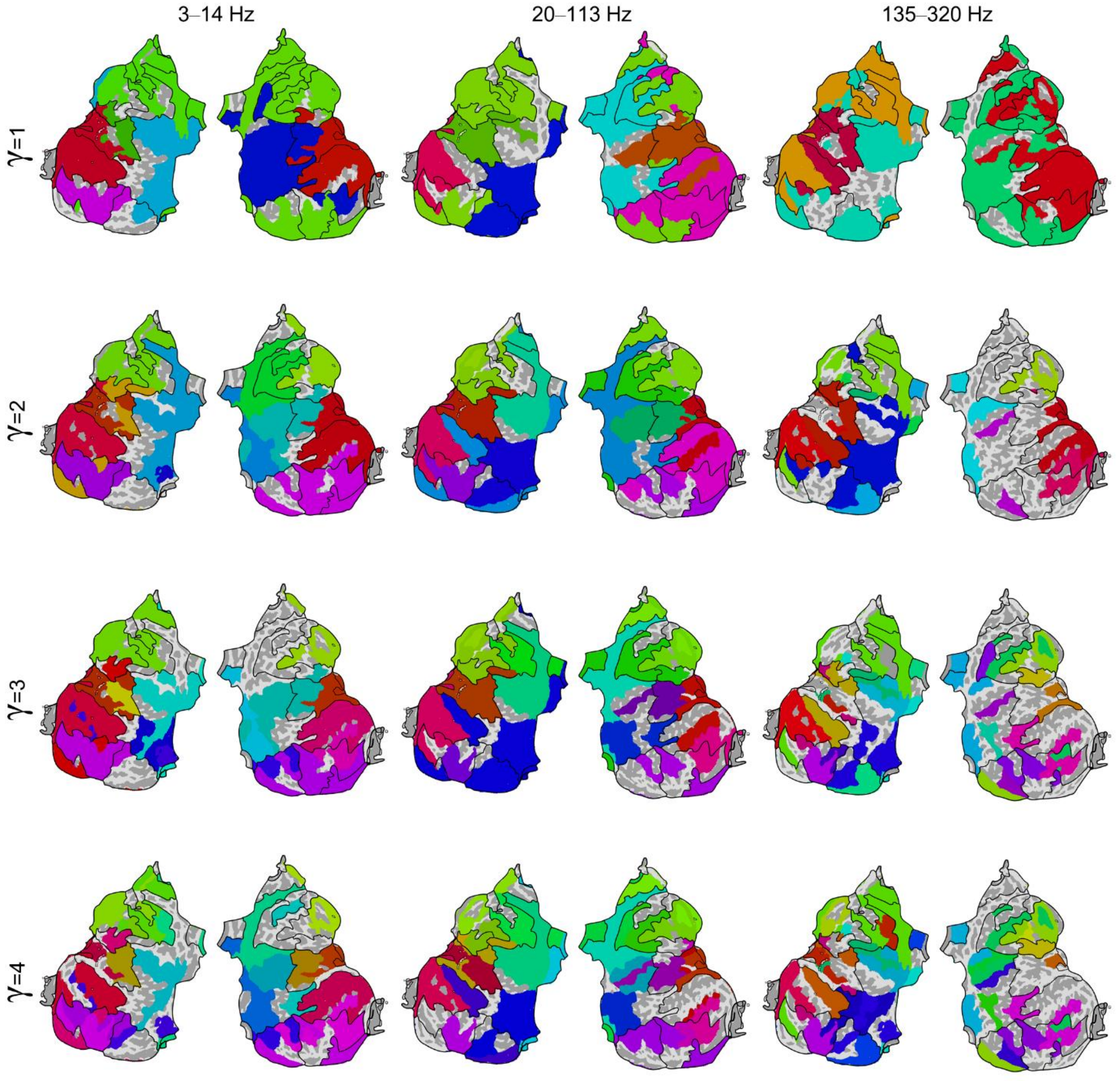


**Figure S13. Modules in connectomes of phase-synchronization generated by retaining top 10 percentile of connections, similar to modules in original connectomes of phase-synchronization.** Flattened cortical surface representations of modules in connectomes of phase-synchronization for 3–14 Hz, 20–113 Hz and 135–320 Hz, at four spatial scales ($\gamma$ = 1 to 4). Connectomes were generated by retaining weights of top 10 percentile of inter-regional connections, while setting all other connections to 0. Black lines on each flattened surface show outlines of consensus modules, *i.e.* sets of regions assigned to the same module across frequencies and spatial scales.

#### 6.6 Robustness of modules to regional amplitude differences

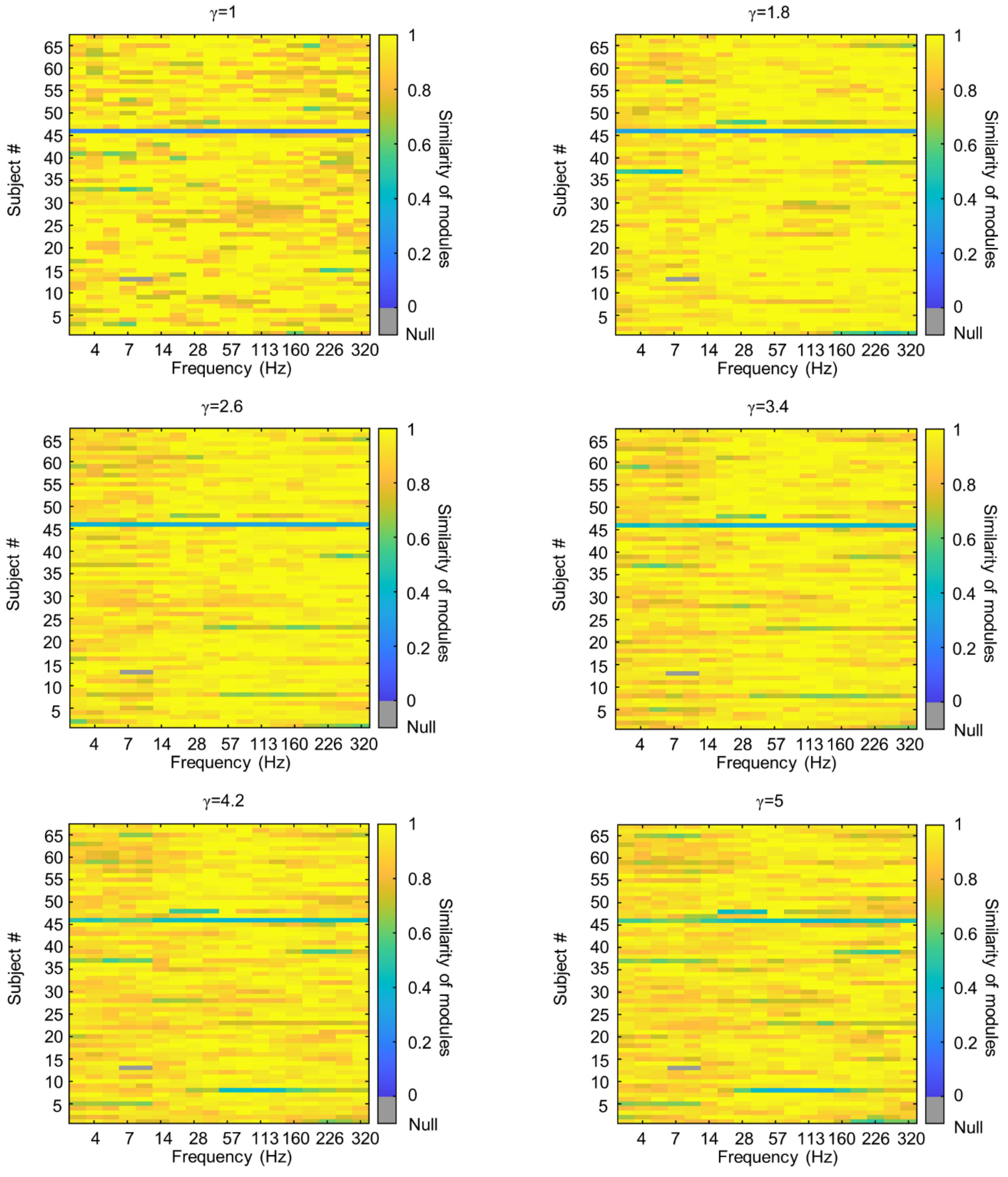


**Figure S14. Amplitude of activity does not confound identification of modules.** Similarity between modules identified on 67 subject-level networks of phase-synchronization across 18 frequencies before and after removing amplitude confound, for six spatial scales (γ = 1, 1.8, 2.6, 3.4, 4.2 and 5). Gray cells represent elements for which we did not assess the amplitude confound.
